# Oncogenic context shapes the fitness landscape of tumor suppression

**DOI:** 10.1101/2022.10.24.511787

**Authors:** Lily M. Blair, Joseph M. Juan, Lafia Sebastian, Vy B. Tran, Wensheng Nie, Gregory D. Wall, Mehmet Gerceker, Ian K. Lai, Edwin A. Apilado, Gabriel Grenot, David Amar, Giorgia Foggetti, Mariana Do Carmo, Zeynep Ugur, Debbie Deng, Alex Chenchik, Maria Paz Zafra, Lukas E. Dow, Katerina Politi, Jonathan J. MacQuitty, Dmitri A. Petrov, Monte M. Winslow, Michael J. Rosen, Ian P. Winters

## Abstract

Tumors acquire alterations in oncogenes and tumor suppressor genes in an adaptive walk through the fitness landscape of tumorigenesis. However, the features of this landscape remain poorly understood and cannot be revealed by human cancer genotyping alone. Here, we use a multiplexed, autochthonous mouse platform to model and quantify the initiation and growth of more than one hundred genotypes of lung tumors across four oncogenic contexts: KRAS G12D, KRAS G12C, BRAF V600E, and EGFR L858R. The resulting fitness landscape is rugged (the effect of tumor suppressor inactivation often switches between beneficial and deleterious depending on the oncogenic context), shows no evidence of diminishing-returns epistasis within variants of the same oncogene, and is inconsistent with expectations of a simple linear signaling relationship among these three oncogenes. Our findings suggest that tumor suppressor effects are strongly context-specific, which limits the set of evolutionary paths that can be taken through the fitness landscape.

## INTRODUCTION

Adaptation by natural selection is the central mechanism of evolution and is at the core of some of the greatest challenges facing humanity: from loss of biodiversity to the spread of infectious disease, to cancer development and resistance to therapy^1-3^. Our capacity to overcome these challenges is dependent on our ability to predict the evolutionary paths taken by such complex and evolving systems. The “fitness landscape”, a map between the genotype and fitness of a biological entity, is a key concept in evolutionary genetics^4^ that provides a framework to understand what kinds of evolutionary paths are possible in a given system.

Theoretical investigations have suggested that fitness landscapes can be broadly categorized as smooth or rugged. In smooth, “Mount Fuji”^5^ landscapes, a mutation that is adaptive in one context will be adaptive in all other contexts. Thus, adaptation is always possible until the fitness peak is reached. In contrast, rugged landscapes, characterized by epistatic interactions wherein the effect of one mutation depends upon others, contain multiple peaks with intervening valleys, inhibiting certain paths. Fitness landscapes can also vary in steepness; individual mutations can be strongly adaptive in steep landscapes while yielding smaller fitness gains in flatter landscapes.

Empirical studies of fitness landscapes have revealed two general observations. First, rugged fitness landscapes consisting of both pairwise and higher-order epistasis are common, making some evolutionary trajectories more probable than others^6^. Second, fixed adaptive mutations reduce the selective advantage of all subsequent mutations—a feature termed diminishing-returns epistasis. Diminishing-returns epistasis was discovered in experimental evolution systems^7-9^, but it remains unknown whether this phenomenon is generalizable across biological systems.

Cancer progression is a quintessential example of a walk on an adaptive fitness landscape, with tumor growth depending on the cooperation of multiple driver mutations^10-12^. While cancer genome sequencing has revealed a vast set of putative cancer drivers—oncogenes and tumor suppressors— mapping the tumor fitness landscape that emerges from coincident alteration of these genes remains a key gap in our understanding of cancer biology. At the level of pairs of drivers, oncogene pairs are known not to have “Mount Fuji”-like additive fitness effects; indeed, the presence of more than one oncogene may even lead to growth arrest or apoptosis^13^. Far less is known about the epistasis of oncogene-tumor suppressor pairs and inferring this dimension of the fitness landscape is the focus of this manuscript.

A common approach to inferring epistatic interactions is to look for non-independence of mutational co-occurrence frequencies in observational cancer genomics data. However, this approach is plagued by two major issues. First, the immense number of genes mutated in human cancers, of which the vast majority are mutated in only a small fraction of tumors, makes the study of mutational co-occurrence statistically underpowered for all but the most frequently mutated genes. Second, even when two driver genes co-occur more or less than expected by chance, this can be due to the confounding biological factors, such as inactivation of a different gene in the same complex, rather than direct functional epistatic interactions^14^. This is especially so in tumors with a high mutational burden where even mutations in known cancer driver genes can be chance passengers, further obscuring the signal. Thus, a global fitness landscape of tumorigenesis cannot be generated from human data alone, and instead requires direct perturbational experiments and functional genomics approaches^14^.

Genetically engineered mouse models are uniquely tractable systems to uncover the phenotypic effects of defined genetic alterations on tumors that develop entirely within their natural *in vivo* microenvironment^15^. Systems that integrate CRISPR/Cas9-mediated somatic genome editing with conventional genetically engineered mouse models of human cancer have increased the scale at which the consequences of tumor suppressor gene inactivation on autochthonous tumorigenesis can be quantified^16, 17^. We recently developed tumor barcoding coupled with high-throughput barcode sequencing (Tuba-seq), which integrates barcoded lentiviral-sgRNA/Cre vectors and barcode sequencing to uncover the number of neoplastic cells in each tumor of each genotype^18, 19^.

Here, we initiate and quantify the development of more than one hundred different genotypes of autochthonous lung tumors. This extensive adaptive fitness landscape overlays inactivation of a broad panel of diverse tumor suppressor genes on top of oncogenic KRAS G12D-, KRAS G12C-, BRAF V600E-, and EGFR L858R-driven lung tumors. *KRAS, EGFR*, and *BRAF* are the three most frequently altered oncogenes in lung adenocarcinoma (LUAD)^20^ and together drive tumorigenesis in over half of patients. Their products are canonically depicted in a linear axis—from EGFR to KRAS to BRAF— within the RAS pathway^20-23^. This linear representation implies that the only dimension upon which these oncogenes can be functionally different is the quantity of downstream MAPK signaling that they drive. However, each of these oncogenes engages additional pathways, which could generate phenotypic differences between these oncogenes in specific contexts^21-23^. Despite the well-established significance of these RAS pathway oncogenes in lung tumorigenesis, it remains unclear the extent to which these off-axis interactions, even if known phenotypically or biochemically, can drive differential *fitness* effects during tumorigenesis. Mutations within *EGFR, KRAS*, and *BRAF* oncogenes are also diverse, and it is unclear whether these mutations are functionally equivalent outside of potential differences in induced RAS signaling.

By generating the most extensive functional survey of oncogene-tumor suppressor interactions to date, we uncover dramatically different tumor suppressive fitness effects across oncogenic contexts, unexpected similarities for oncogenes with strong differences in tumor-driving potential, and surprising effects of off-axis signaling.

## RESULTS

### Oncogenic KRAS G12C is less potent than KRAS G12D in driving lung tumorigenesis

Oncogenic KRAS mutations, predominantly within codon 12, occur in ∼25% of human LUAD (**Supplementary Table 1**)^24, 25^. To enable CRISPR/Cas9-mediated somatic genome editing in the context of different oncogenic KRAS variants, we generated mice with Cre/lox-regulated alleles of KRAS G12C (*Kras*^*LSL-G12C*^)^26^ or KRAS G12D (*Kras*^*LSL-G12D*^)^27^ and a Cre/lox-regulated Cas9 allele (*H11*^*LSL-Cas9*^; **Fig. 1a**)^28^. While the impact of inactivating diverse tumor suppressor genes on KRAS G12D-driven lung cancer growth has been investigated previously^19, 29^, the functional landscape of tumor suppression within KRAS G12C-driven lung cancer *in vivo* remains entirely uncharacterized, even though KRAS G12C is the most common oncogenic KRAS variant in human lung cancer (**Supplementary Fig. 1**). To broadly uncover the genetic interactions between tumor suppressor genes and these oncogenic KRAS variants *in vivo*, we generated barcoded Lenti-sgRNA/Cre vectors targeting 28 known and putative tumor suppressor genes that are recurrently mutated in human LUAD and represent key cancer pathways (**Fig. 1a, Supplementary Table 2**, and **Methods)**^20, 25, 30^. We generated a pool of barcoded Lenti-sgRNA/Cre vectors, which included vectors targeting each of these genes as well as control Lenti-sgRNA/Cre vectors with non-targeting sgRNAs (sg*NT*) and active-cutting sgRNAs (sg*AC*) that target an inert region of the genome (*Rosa26*; sg*R26*) (Lenti-D2G^28-Pool^/Cre; **Fig. 1a** and **Supplementary Fig. 2a**).

**Figure 1.**
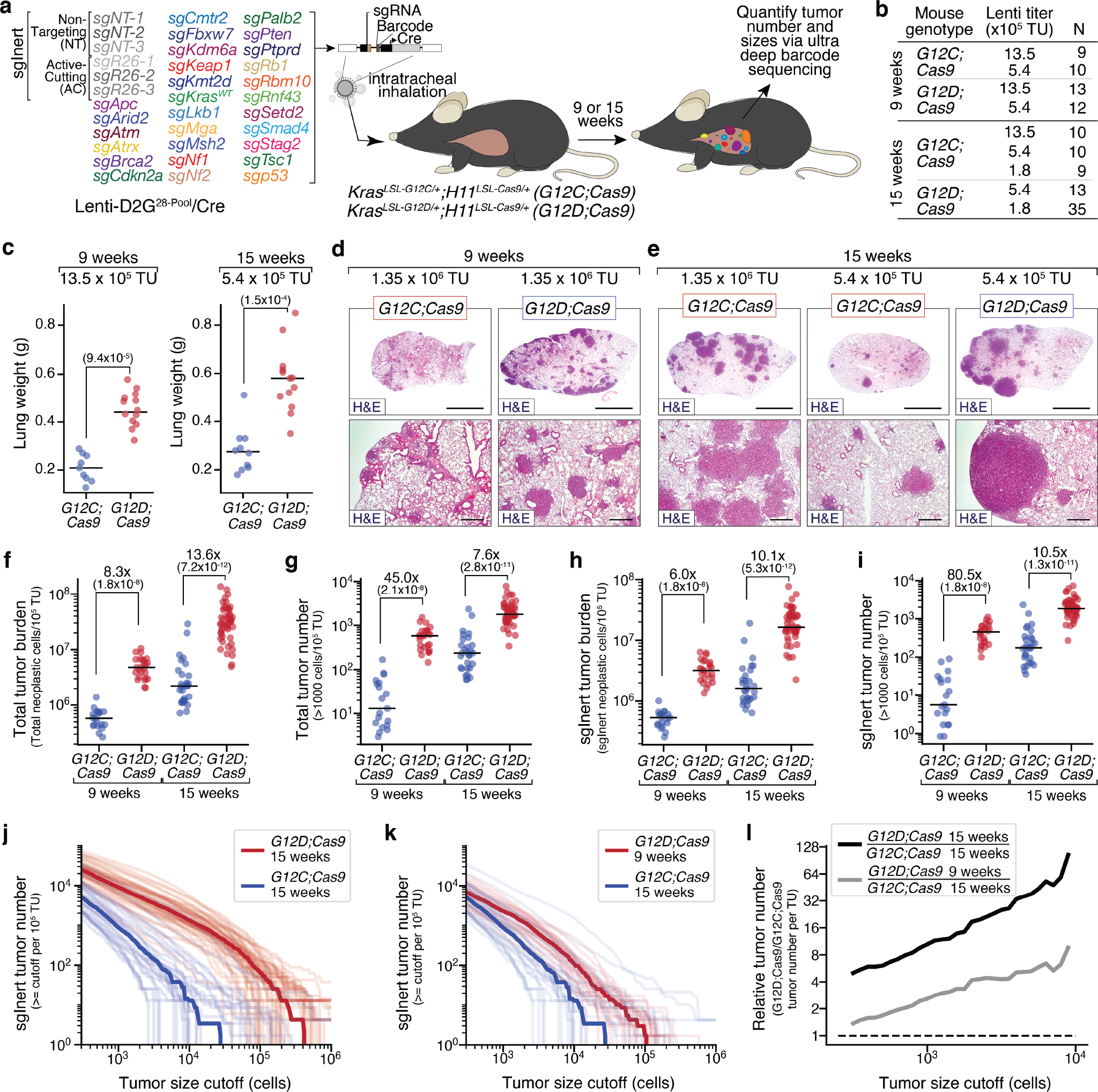
Oncogenic KRAS G12C has reduced ability to drive initiation and growth of lung tumors *in vivo* relative to oncogenic KRAS G12D. **a**. Experimental schematic depicting the composition of the pool of barcoded Lenti-sgRNA/Cre vectors (Lenti-D2G^28-Pool^/Cre), mouse genotypes, analysis time points, and readouts. b.Genotype, time point, lentiviral titer, and number of mice in each group. c.Lung weights of mice transduced with the indicated titers of Lenti-D2G^28-Pool^/Cre. Genotype and time post-tumor initiation are indicated. Each dot represents a mouse and the bar is the median. Fold difference between medians and significance calculated using a wilcoxon rank sum test (p-values < number in parentheses) are shown. **d**,**e**. Representative histology of lungs from mice. Mouse genotype, virus titer delivered to each mouse, and time post-tumor initiation are shown. Top scale bars = 3 mm; bottom scale bars = 500 µM. **f-i**. Total number of neoplastic cells (**f**) and total number of tumors greater than 1000 cells in size (**g**) across all Lenti-sgRNA/Cre vectors, normalized to viral titer. Total number of neoplastic tumors cells (**h**) and total number of tumors greater than 1000 cells in size (**i**) only for Lenti-sgInert/Cre vectors (tumors driven by oncogenic Kras alone), normalized to viral titer. Mouse genotypes and time points are indicated. Each dot represents a mouse and the bar is the median. Fold difference and significance calculated using a wilcoxon rank sum test (p-values < number in parentheses) are shown. **j**,**k**. Number of tumors at or above the tumor size cutoff in *G12D;Cas9* mice at 15 weeks and *G12C;Cas9* mice at 15 weeks (**j**) or *G12D;Cas9* at 9 weeks and *G12C;Cas9* at 15 weeks post-tumor initiation (**k**). Each transparent line represents a mouse and the solid line is the median tumor number. **l**. Fold-change in median tumor number between *G12D;Cas9* and *G12C;Cas9* at 15 weeks (black line) and *G12D;Cas9* at 9 weeks versus *G12C;Cas9* at 15 weeks (gray line) post-tumor initiation.

We initiated tumors by delivering Lenti-D2G^28-Pool^/Cre to intratracheally intubated *Kras*^*LSL-G12C*^*;H11*^*LSL-Cas9*^ (*G12C;Cas9*) and *Kras*^*LSL-G12D*^*;H11*^*LSL-Cas9*^ (*G12D;Cas9*) mice (**Fig. 1a**). Given the uncertain oncogenicity of KRAS G12C relative to KRAS G12D, we initiated tumors with several different titers of Lenti-D2G^28-Pool^/Cre (from 1.8×10^5^ to 1.35×10^6^ TU (transduction units)/mouse) and analyzed cohorts of mice at 9 and 15 weeks post-tumor initiation (**Fig. 1b**; N = between 9 and 35 mice per group). At the time of lung collection, *G12C;Cas9* mice had noticeably fewer and smaller surface lung tumors and significantly lower lung weights relative to titer- and timepoint-matched *G12D;Cas9* mice (**Fig. 1c** and **Supplementary Fig. 3a,b**). Histology indicated that *G12C;Cas9* mice had fewer tumors and these appeared smaller than those in *G12D;Cas9* mice (**Fig. 1d,e**). Lung tumors in both backgrounds were hyperplasias, adenomas, and adenocarcinomas. These results suggest that KRAS G12C is less potent than KRAS G12D in driving lung tumorigenesis, consistent with previous studies using *in vivo* models of lung and pancreatic cancer^26, 31^.

### KRAS G12C induces fewer and smaller tumors than KRAS G12D

To quantify the number and size of the tumors in each mouse, and to better understand the dynamics of lung tumor growth driven by these different oncogenic KRAS variants, we performed Tuba-seq on DNA extracted from bulk tumor-bearing lungs from mice across the different titers and timepoints (**Supplementary Fig. 2b**). Tuba-seq accurately quantifies the number of neoplastic cells in each tumor (cells directly descending from the tumor cell of origin) through deep sequencing of the multi-component barcode encoded within each genomically-integrated lentivirus (**Supplementary Fig. 2a-c**)^19, 32^. This allowed us to estimate total tumor burden (total neoplastic cells/TU) and total tumor number (number of clonal barcoded tumors with >1000 neoplastic cells/TU) in each mouse (**Supplementary Fig. 2a-c** and **Methods**)^19, 32^. 9 weeks post-tumor initiation, total tumor burden (normalized to titer) was >8-fold lower in *G12C;Cas9* mice than in *G12D;Cas9* mice (**Fig. 1f** and **Supplementary Fig. 3 b,c**; p<2×10^−8^). This difference was slightly greater and still highly significant 15 weeks post-tumor initiation (**Fig. 1f** and **Supplementary Fig. 3 b,c**; p<7×10^−12^). At both 9 and 15 weeks post-tumor initiation, *G12C;Cas9* mice also had many fewer tumors per TU than *G12D;Cas9* mice (**Fig. 1g** and **Supplementary Fig. 3 b,c**; p<2×10^−8^). Tumor number increased linearly with titer, consistent with a lack of inter-tumor competition even at high tumor burden. Thus, when considering all tumors independently of their engineered tumor suppressor inactivation, KRAS G12C drives substantially less neoplastic growth than KRAS G12D.

Lenti-sg*NT*/Cre and Lenti-sg*R26*/Cre (sgInert) vectors induce the expression of oncogenic KRAS from the engineered alleles without CRISPR/Cas9-mediated inactivation of any gene, generating tumors driven solely by oncogenic KRAS. Before exploring the impact of inactivation of each tumor suppressor gene on tumor growth, we restricted our analysis to these sgInert “KRAS-only” tumors. sgInert tumor burden and tumor number were also dramatically lower in *G12C;Cas9* mice relative to *G12D;Cas9* mice, at both 9 and 15 weeks post-tumor initiation (**Fig. 1h,i** and **Supplementary Fig. 3b-d**; all p<10^−**7**^).

To further investigate the different abilities of oncogenic KRAS G12C and KRAS G12D to initiate lung tumors and drive their growth, we explored the distribution of sgInert tumor sizes (**Fig. 1j-l** and **Supplementary Fig. 3e-g**). Comparison of the two models 15 weeks post-tumor initiation revealed fewer KRAS G12C tumors than KRAS G12D tumors above any minimum size cutoff (**Fig. 1j**). Furthermore, the KRAS G12D tumor size distribution had a longer tail of large tumors, suggesting that its increased tumor number might be driven by more rapid growth than tumors driven by KRAS G12C (**Fig. 1j**). In support of this notion, the shape of the KRAS G12C tumor size distribution 15 weeks post-tumor initiation was similar to that of the KRAS G12D tumor size distribution at the earlier 9-week timepoint (**Fig. 1k**). However, while the shapes of the distributions at these two timepoints were quite well matched (**Fig. 1k**), KRAS G12D consistently produced ∼2-4x greater tumor number than KRAS G12C, suggesting that KRAS G12D may also drive greater levels of tumor initiation (**Fig. 1l**). These results are all consistent with a model in which KRAS G12C is less potent at initiating lung tumors and less able to drive the expansion of established tumors *in vivo* than KRAS G12D.

### Diverse tumor suppressor genes have strikingly similar effects on the initiation and growth of KRAS G12C- and KRAS G12D-driven lung tumors

Having used Tuba-seq to uncover differences in the baseline ability of KRAS G12C and KRAS G12D to initiate lung tumors and drive their growth, we next analyzed the impact of inactivating each of the 28 putative tumor suppressor genes on the growth of lung tumors driven by these oncogenes (**Fig. 2a,b** and **Supplementary Fig. 4a**). To compare the effects of inactivating each targeted gene across oncogenes, we analyzed all tumors above oncogene-specific tumor size (number of neoplastic cells) cutoffs that matched the number of sgInert tumors in each oncogenic context (**Methods**). Matching cutoffs in this way allowed us to account for differential oncogene-intrinsic growth dynamics. We used a minimum tumor size cutoff of 1600 cells for *G12D;Cas9* mice at 15 weeks post-tumor initiation, 600 cells for *G12D;Cas9* mice at 9 weeks, 400 cells for *G12C;Cas9* mice at 15 weeks, and 300 cells for *G12C;Cas9* at 9 weeks. We then compared the sizes of tumors in which each tumor suppressor gene was targeted to the sizes of sgInert tumors.

**Figure 2.**
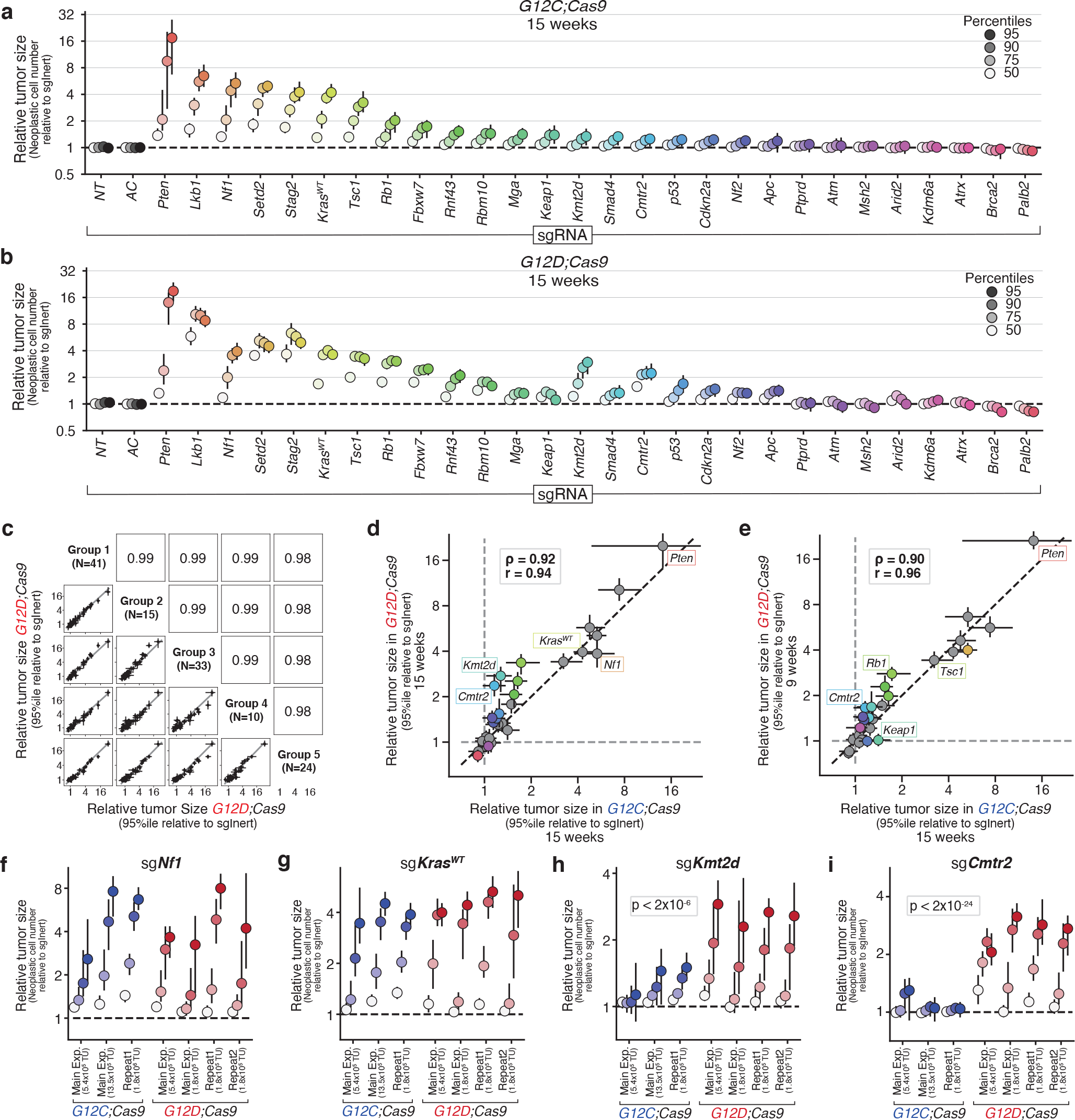
Tumor suppressor genes have strikingly similar effects on the initiation and growth of KRAS G12C- and G12D-driven lung tumors. **a,b**. Relative size (neoplastic cells) of the tumor at the indicated percentiles of the tumor size distributions for barcoded Lenti-sgRNA/Cre vectors targeting each gene, relative to the size of the sgInert tumor at the same percentile, in *G12C;Cas9* mice (**a**) and *G12D;Cas9* mice (**b**) at 15 weeks after tumor initiation. 95% confidence intervals are shown. **c**. 95^th^ percentile relative tumor sizes (relative to sgInert) for 5 of the *G12D;Cas9* study groups (See Supplementary Fig 5a,b for comparisons between addditional study groups). Each point represents the tumors initiated with one Lenti-sgRNA/Cre vector and the bars are the 95^th^ percent confidence intervals. Grey line indicates equal effect. Pearson r is indicated. **d,e**. Relative size of the tumor at the 95^th^ percentile of the tumor size distributions in *G12D;Cas9* mice at 15 weeks (**d**) or *G12D;Cas9* mice at 9 weeks (**e**) versus in *G12C;Cas9* mice at 15 weeks after tumor initiation. Each dot represents the tumors initiated from one Lenti-sgRNA/Cre vector and the bars are the 95^th^ percent confidence intervals. Genes where the 95% CI excluded no effect in *G12C;Cas9* and *G12D;Cas9* mice are shown in color and some key genes are labeled. The black dotted line indicates equal effect. Spearman rank-order correlation (ρ) and Pearson correlation (r) are indicated. **f-i**. Relative size of the tumor at the indicated percentiles (see legend in **a,b**) of the tumor size distributions for barcoded Lenti-sgRNA/Cre vectors targeting *Nf1* (**f**), *Kras*^*WT*^ (**g**), *Kmt2d* (**h**), and *Cmtr2* (**i**) across multiple arms of our main experiment and repeat studies in *G12C;Cas9* and *G12D;Cas9* mice. Significance of oncogene differences at the 95^th^ percentile is calculated by combining the study groups for each oncogene using inverse variance weighting and comparing the resulting means and variances under a normally distributed null. Bonferroni-corrected p-values shown for significant genes.

Inactivation of many of these genes led to the development of larger KRAS G12C- and KRAS G12D-driven tumors in our study (**Fig. 2a,b**). These tumor suppressive effects were highly reproducible across eleven *G12D;Cas9* study groups (pre-defined cohorts of mice of identical genotype, administered viral titer, date of tumor initiation, and date of take down) and four *G12C;Cas9* study groups (**Fig. 2c** and **Supplementary Fig. 5**; data from 243 *G12D;Cas9* mice and 47 *G12C;Cas9* mice; Pearson r≥0.95 and r≥0.87, respectively, for each comparison).

Comparing size distributions of tumors of each tumor suppressor genotype revealed consistent effects between KRAS G12C- and KRAS G12D-driven lung tumors. Despite the differences in oncogenic potential of KRAS G12C and KRAS G12D, the tumor suppressive effects at 15 weeks post-initiation were highly correlated (**Fig. 2a,b,d**; Spearman ρ=0.92). Tumor suppressive effects were also highly correlated between KRAS G12C-driven tumors at 15 weeks post-tumor initiation and KRAS G12D-driven tumors at 9 weeks post-tumor initiation, when sgInert tumors were most similar (**Fig. 2e**; Spearman ρ=0.90). In fact, across all comparisons of timepoints and oncogenic alleles, tumor suppressive effects were well correlated (**Fig. 2d,e** and **Supplementary Fig. 6**; all Spearman and Pearson correlations ρ≥0.88).

Our pool contained Lenti-sgRNA/Cre vectors targeting two established negative regulators of oncogenic KRAS signaling: the NF1 RAS GTPase-activating protein and wild type KRAS (*Kras*^*WT*^). Inactivation of either *Nf1* or *Kras*^*WT*^ has been shown to increase downstream signaling and enable faster growth of oncogenic KRAS G12D-driven lung tumors *in vivo*^19, 33, 34^. Contrary to our expectation that inactivation of these negative regulators would have a greater effect on tumors driven by the weaker KRAS G12C variant, inactivation of *Nf1* or *Kras*^*WT*^ increased KRAS G12C- and KRAS G12D-driven tumor growth to the same extent. This result was consistent across study groups in multiple independent studies (**Fig. 2f,g**). Thus, despite the large difference in the ability of KRAS G12C and KRAS G12D to drive tumor growth, as well as their known biochemical differences^35^, lung tumor growth driven by these oncogenic variants was similarly affected by alterations in oncogene-proximal tumor suppressors.

While the impact of inactivating most tumor suppressor genes was similar in lung tumors driven by KRAS G12C and KRAS G12D, there were some notable exceptions. Inactivation of the H3K4 mono- and di-methyltransferase KMT2D or the cap-specific mRNA methyltransferase CMTR2 increased the growth of KRAS G12C-driven tumors less than KRAS G12D-driven tumors (**Fig. 2h,i**; p<2×10^−6^, 2×10^−24^, respectively). These results are inconsistent with the model of diminishing returns epistasis where adaptive mutations are expected to provide greater fitness benefit on the less fit genetic background—in this case, KRAS G12C. These differences were again consistent across different studies, titers, and timepoints post-tumor initiation (**Fig. 2h,i**).

### Oncogenic BRAF and EGFR have distinct tumor-initiating and growth-promoting abilities

Oncogenic mutations in BRAF and EGFR occur frequently in human LUAD, underscoring the importance of activation of the RAS pathway in a large fraction of these tumors (**Supplementary Table 1**)^20, 25^. Oncogenic BRAF mutations (including those at the hotspot V600) occur in ∼6% of LUAD, while oncogenic EGFR mutations occur in ∼27% (AACR Project GENIE) of LUAD (**Supplementary Table 1**)^20, 36^. BRAF- and EGFR-driven lung cancers have been modelled in mice using a Cre/lox-regulated conditionally activatable allele of BRAF V600E (*Braf*^*CA-V600E*^)^37^ and a doxycycline regulated *EGFR*^*L858R*^ transgene^38, 39^. To quantify the ability of oncogenic BRAF and EGFR to initiate lung tumors and drive their expansion *in vivo*, as well as to uncover whether tumor suppressor effects are consistent across these oncogenic contexts in lung cancer, we initiated tumors with Lenti-D2G^28-Pool^/Cre in *Braf*^*CA-V600E*^*;H11*^*LSL-Cas9*^ (*Braf;Cas9*) and *tetO-EGFR*^*L858R*^*;Rosa26*^*LSL-rtTA3-ires-mKate/LSL-Cas9-2a-GFP*^ (*Egfr;Cas9*) mice (**Fig. 3a**)^38^. Mice received several different titers of Lenti-D2G^28-Pool^/Cre (from 1.6×10^4^ to 5×10^6^ TU/mouse) and were analyzed 15 weeks post-tumor initiation (**Fig. 3b**). Consistent with previous observations, *Braf;Cas9* mice developed lung tumors that appeared more uniform in size than oncogenic KRAS-driven or oncogenic EGFR-driven tumors (**Fig. 1d,e** and **Fig. 3c**. Lung weights shown in **Supplementary Fig. 7a,b**)^37, 40^. BRAF and EGFR-driven tumors were adenomas and adenocarcinomas (**Fig. 3c**).

**Figure 3.**
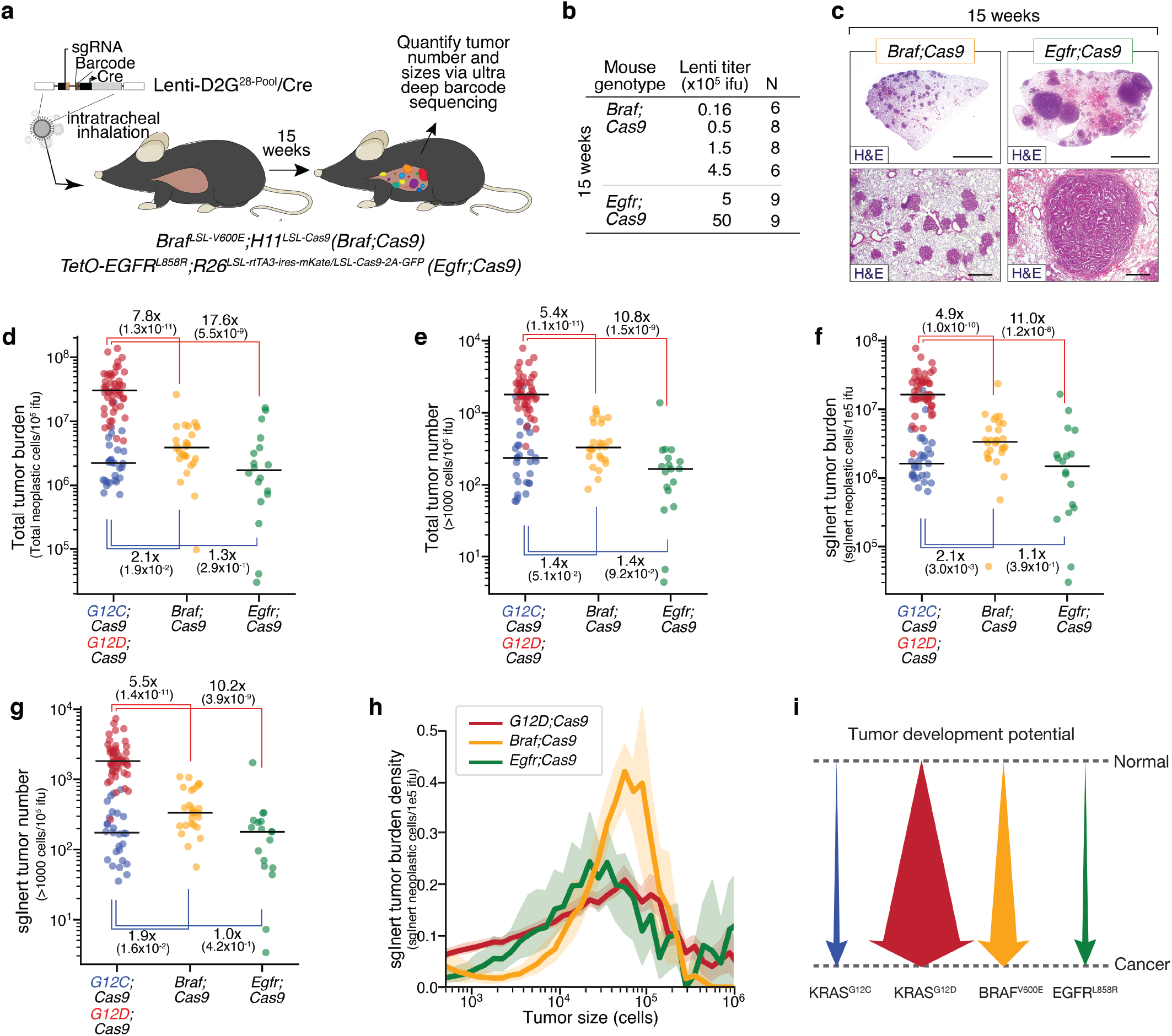
Oncogenic BRAF, EGFR and KRAS have different abilities to initiate lung tumorigenesis and drive tumor growth. **a.** Experimental schematic showing the design of barcoded Lenti-sgRNA/Cre vectors (Lenti-D2G^28-Pool^/Cre), mouse genotypes, analysis timepoints. **b.** Mouse genotype, time-point, lentiviral titer, and number of mice in each experimental group. **c.** Representative histology of lungs from mice. Mouse genotype, viral titer, and time point after tumor initiation are shown. Top scale bars = 3mm; bottom scale bars = 500 µM. Titer represented in the *Braf;Cas9* image was 900,000 TU, from a mouse in a separate titering experiment that used a similar virus pool. Titer represented in the *Egfr;Cas9* image was 5,000,000 TU. **d-g**. Total number of neoplastic cells (**d**) and total number of tumors greater than 1000 cells in size (**e**) across all Lenti-sgRNA/Cre vectors, normalized to viral titer. Total number of neoplastic tumors cells (**f**) and total number of tumors greater than 1000 cells in size (**g**) only for Lenti-sgInert/Cre vectors (tumors driven by oncogene alone), normalized to viral titer. Mouse genotypes are indicated. Each dot represents a mouse and the bar is the median. Fold differences between medians and significance calculated using a wilcoxon rank sum test (p-values < number in parentheses) are shown. Fold differences are ratios of the following pairs, moving clockwise from the upper left: *G12D;Cas9 / Braf;Cas9, G12D;Cas9 / Egfr;Cas9, G12C;Cas9 / Egfr;Cas9, Braf;Cas9 / G12C;Cas9*. **h**. The density function of sgInert tumor burden as a function of log(Tumor size) 9 weeks for *G12D;Cas9, Braf;Cas9*, and *Egfr;Cas9*. Comparison to *G12C;Cas9* can be found in **Supplementary Fig. 3f,g.** **i**. Schematic representation of the ability of each indicated oncogenic allele to drive *in vivo* lung tumor formation.

Tuba-seq analysis of DNA extracted from bulk tumor-bearing lungs allowed us to quantify tumor burden and size across mouse genotypes, and thus determine the tumorigenic potential of BRAF V600E and EGFR L858R relative to KRAS G12C and KRAS G12D. Total tumor burden and total tumor number in *Braf;Cas9* mice was higher than in *G12C;Cas9* mice but >5-fold lower than in *G12D;Cas9* mice (**Fig. 3d,e**; p<1.9^-2^, p<1.3^-11^). *Egfr;Cas9* mice had slightly lower tumor burden and number than *Braf;Cas9* mice and >10-fold fewer tumors than *G12D;Cas9* mice (**Fig. 3d,e**; p<1.5^-9^). These results remained consistent after restricting the analysis to sgInert-containing tumors driven by oncogenic BRAF or EGFR (**Fig. 3f,g**; p<1.6^-2^).

Interestingly, the distribution of BRAF-driven tumor sizes was strikingly different from that of other oncogenic contexts. Exceptionally large tumors accounted for a much smaller percentage of the total burden of neoplastic tumor cells in *Braf;Cas9* mice than in any other mouse genotype: only 0.9% of neoplastic cells in *Braf;Cas9* mice were from tumors larger than 300,000 cells compared with 13.1% and 25.4% for *G12D;Cas9* and *Egfr;Cas9* mice, respectively—consistent with previous reports that BRAF tumors hit a maximum size threshold and stop growing^40^. However, unlike in *G12C;Cas9, G12D;Cas9*, and *Egfr;Cas9* mice, a majority of the total neoplastic burden in *Braf;Cas9* mice arose from tumors in the 30,000 to 300,000 cell range, the order of magnitude just below the largest tumors: 62.2% in *Braf;Cas9* mice compared with 36.0% and 32.3% in *G12D;Cas9* and *Egfr;Cas9* mice, respectively (**Fig. 3h** and **Supplementary Fig. 7c,d**). In comparison to *Braf;Cas9* mice, a much greater fraction of the total neoplastic burden in *G12D;Cas9* mice arose from smaller tumors. The difficulty in quantifying these smaller tumors using histological methods might explain why previous studies have suggested that BRAF V600E is a stronger driver of lung tumorigenesis than KRAS G12D^41, 42^. Collectively, these results indicate that different oncogenes in the EGFR/KRAS/BRAF axis have dramatically different effects on tumor initiation and growth (**Fig. 3i**).

### Oncogenic BRAF and EGFR redefine the landscape of tumor growth suppression

We next investigated the impact of tumor suppressor gene inactivation on the growth of BRAF V600E- and EGFR L858R-driven lung tumors and compared tumor suppressor effects across all four oncogenic alleles. Very few coincident tumor suppressor alterations have been investigated in the context of oncogenic BRAF-driven autochthonous lung tumors, and the extent to which tumor suppressor effects differ in EGFR-driven tumors remains poorly understood (**Supplementary Fig. 1**)^38, 43-46^. Interestingly, the overall tumor suppressive landscapes of BRAF- and EGFR-driven lung tumors were dramatically different from each other as well as from oncogenic KRAS-driven tumors (**Fig. 2a,b** and **Fig. 4a,b**). Indeed, the effects of tumor suppressor inactivation on growth of across oncogenic KRAS-, BRAF-, and EGFR-driven tumors were uncorrelated (**Fig. 4c,d**; Spearman ρ=0.41 for BRAF versus G12C, ρ=0.14 for EGFR versus G12C). While inactivation of some tumor suppressor genes increased growth across all contexts (e.g., *Pten*), those were the exception (**Fig. 2, Fig. 4**, and **Supplementary Fig. 8**). Importantly, several tumor suppressor genes impacted tumorigenesis as anticipated. Inactivation of *Nf1* increased size of KRAS and EGFR-driven tumors while having no effect on BRAF-driven tumors (**Supplementary Fig. 8d**). Inactivation of *Kras*^*WT*^ increased the growth of KRAS G12C- and KRAS G12D-driven tumors but had no effect of BRAF-driven tumors and reduced the growth of EGFR-driven tumors (consistent with KRAS being an important downstream effector) (**Fig. 4e**).

**Figure 4.**
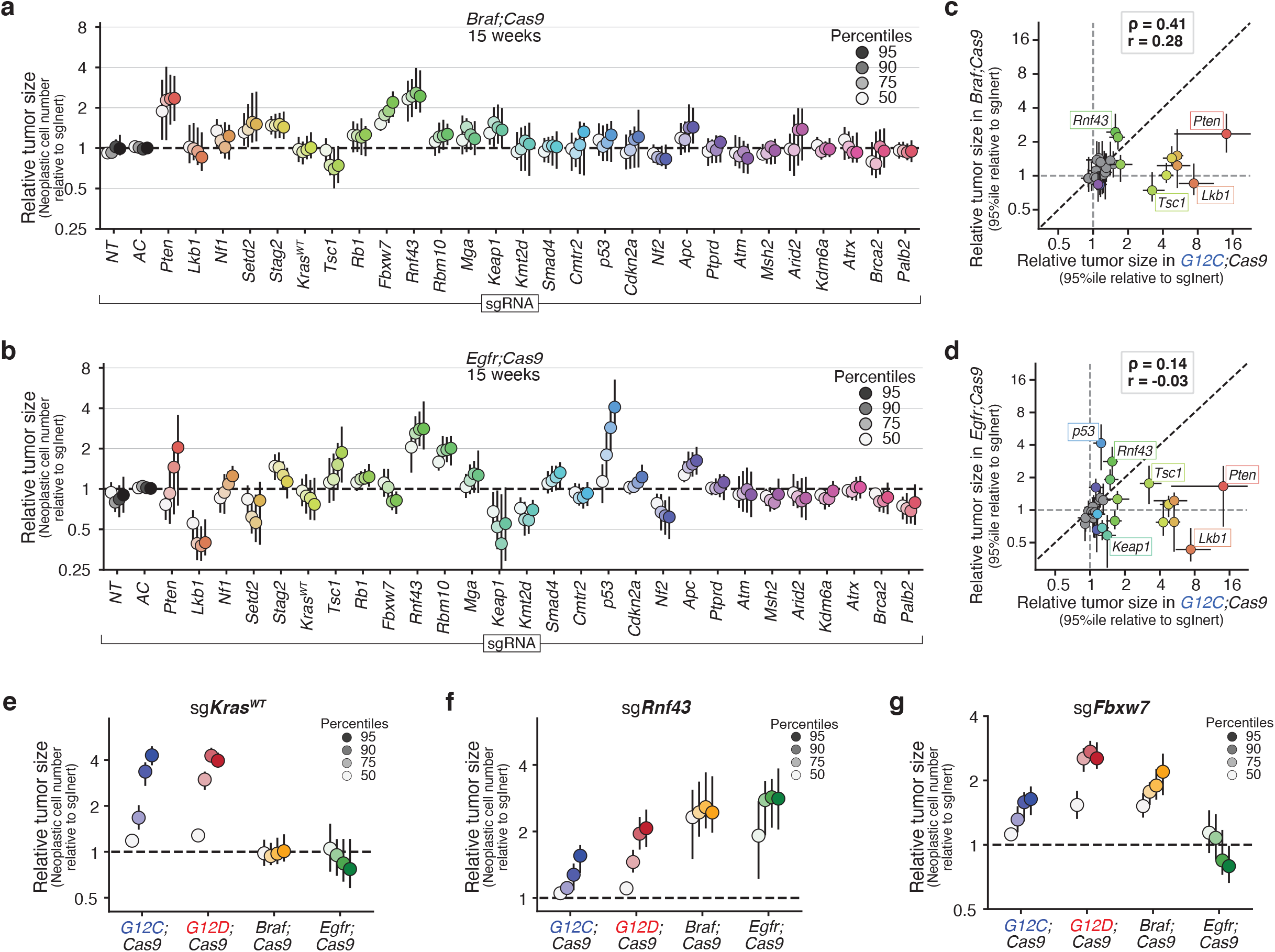
Oncogenic driver defines the landscape of tumor growth suppresion in lung cancer. **a,b**. Relative size (neoplastic cells) of the tumor at the indicated percentiles of the tumor size distributions for barcoded Lenti-sgRNA/Cre vectors targeting each gene, relative to the size of the sgInert tumor at the same percentile, in *Braf;Cas9* mice (**a**) and *Egfr;Cas9* mice (**b**) at 15 weeks after tumor initiation. **c,d**. Relative size of the tumor at the 95^th^ percentile of the tumor size distributions in *G12C;Cas9* versus *Braf;Cas9* mice (**c**) and *Egfr;Cas9* mice (**d**). Each dot represents the tumors initiated from one Lenti-sgRNA/Cre vector and the bars are the 95^th^ percent confidence intervals. Genes where the 95% CI excluded no effect are shown in color and some key genes are labeled. Black dotted line indicates equal effect. Spearman rank-order correlation (ρ) and Pearson correlation (r) are indicated. **e-g**. Relative size of the tumor at the indicated percentiles of the tumor size distributions for Lenti-sgRNA/Cre vectors targeting *Kras*^*WT*^ (**e**), *Rnf43* (**f**), and *Fbxw7* (**g**) and in tumors in the indicated genotypes of mice.

Inactivation of many tumor suppressor genes had strikingly different effects on the growth of BRAF-driven tumors compared to tumors driven by either KRAS variant (**Fig. 4c** and **Supplementary Fig. 8a**). While the tumor suppressive effects of inactivating *Pten, Rnf43*, and *Apc* are consistent with previous data on these genes and/or related pathways in BRAF-driven lung cancer^40, 43-45, 47-50^, the general decreases in magnitude relative to oncogenic KRAS were unexpected (**Fig. 4f** and **Supplementary Fig. 8e**). The effect of coincident tumor suppressor inactivation could generally be reduced due to diminishing-returns epistasis in fast growing BRAF V600E-driven tumors. However, lower effect magnitudes were not universal as inactivation of *Rnf43* or *Fbxw7* increased tumor growth as much or more in *Braf;Cas9* mice than in *G12C;Cas9* mice (**Fig. 4f,g**). Thus, the impact of certain tumor suppressor pathways on tumor growth largely depends on which oncogene is activated.

The differences between tumor suppressive effects in EGFR-driven lung cancer and the other oncogenic contexts were even more pronounced (**Fig. 4b,d**). Inactivation of many genes that were functional tumor suppressors in KRAS-driven lung tumors, including *Lkb1, Setd2*, and *Kmt2d*, were deleterious in EGFR-driven lung tumors (**Fig. 4b,d** and **Supplementary Fig. 8b,g-i**). Conversely, inactivation of *p53* increased the overall growth of EGFR-driven lung tumors more than in any other oncogenic context (**Supplementary Fig. 8f**). These differences represent the clearest indication of a rugged landscape of oncogene-tumor suppressor interactions; whether a second step (tumor suppressor inactivation) led uphill or downhill depended strongly on which first uphill step was taken (EGFR vs KRAS or BRAF).

### Genetic interactions between oncogenes and tumor suppressors impact the earliest stages of tumor development

The initiation of tumors using the exact same virus pool in mice with and without the *Cas9* allele enables quantification of the impact of each gene on tumor number, as the number of tumors in mice without the *Cas9* allele defines the representation of each virus in the pool (**Supplementary Fig. 9**)^19^. As anticipated, mice with conditional oncogene alleles but lacking the *H11*^*LSL-Cas9*^ allele (Cas9-negative mice) transduced with Lenti-D2G^28-Pool^/Cre had much lower overall tumor burden than their Cas9-positive counterparts, and no Lenti-sgRNA/Cre vectors had any effect on tumor sizes (**Supplementary Fig. 4c**). Within each oncogenic context, we assessed parallel cohorts of Cas9-negative mice, allowing us to quantify the impact of each tumor suppressor gene on tumor initiation/early tumor expansion (**Fig. 5a-d** and **Supplementary Fig. 9b-e**). Given the resolution of our approach, we note that tumor number measurements represent the number of tumors with >500 neoplastic cells.

**Figure 5.**
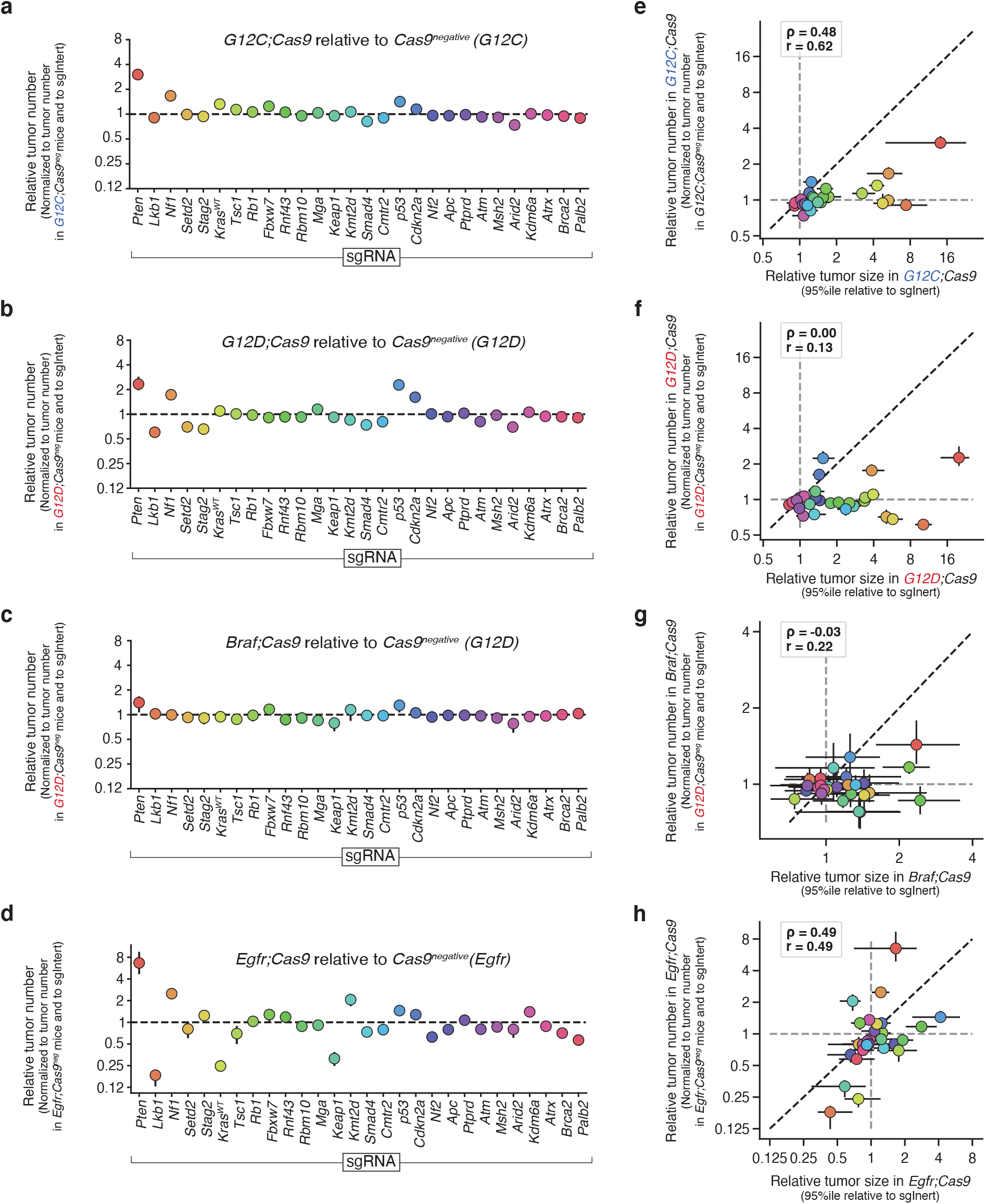
The impact of different tumor suppressors on lung tumor number is dependent on oncogenic context and largely independent of effects on tumor growth. **a-d**. Impact of inactivating each gene on relative tumor number in the indicated genotypes of mice. **e-h**. Relative size of the tumor at the 95^th^ percentile of the tumor size distributions versus relative tumor number in the indicated genotypes of mice. The impact of inactivation tumor suppressor genes on tumor number enrichment and tumor size are not correlated. Each dot represents the tumors initiated from one Lenti-sgRNA/Cre vector in the context of the oncogene indicated on the x- and y-axes.

As was the case for tumor growth effects, inactivation of many genes had similar effects on tumor initiation of KRAS G12C- and KRAS G12D-driven tumors (**Fig. 5a,b** and **Supplementary Fig. 10a**). However, the impact of different tumor suppressors on tumor number varied across oncogenic KRAS, BRAF, and EGFR contexts (**Fig. 5** and **Supplementary Fig. 10**). Interestingly, in *Braf;Cas9* mice, inactivation of tumor suppressors had little effect on tumor number (**Fig. 5c**). Conversely, the number of EGFR-driven tumors was greatly impacted by coincident tumor suppressor inactivation. These effects were large in magnitude (e.g., >6-fold increase for sg*Pten*) and included many genes that reduce tumor number (e.g., >4-fold decrease for sg*Lkb1*), suggesting several of these tumor suppressor genes do not in fact suppress EGFR-driven tumors (**Fig. 5d**). Thus, much like the effects on tumor growth, tumor initiation/early expansion is highly context-dependent with complex and diverse genetic interactions influencing even the earliest steps of lung carcinogenesis.

Finally, as was found before in the KRAS-G12D context, across all four oncogenic contexts in our study the impact of inactivating tumor suppressor genes on tumor initiation/early expansion and tumor growth did not correlate (**Fig. 5e-h)**. This suggests that the genes and pathways that regulate the earliest stages of tumorigenesis are largely non-overlapping with those that modulate later tumor growth.

### *In vivo* tumor suppressive effects positively correlate with the frequency of tumor suppressor alterations in human tumors when the burden of passenger mutations is low

Finally, we performed a retrospective analysis of whether the frequencies of tumor suppressor alterations in 2,204 patients with LUAD and tumor genomic profiles (AACR Project GENIE) correlate with the fitness effects elucidated using our *in vivo* models. Such a correlation is expected if the mutation frequencies are driven by effects on tumor growth that our model recapitulates, but may be undermined by the fact that i) tumor suppressor genes could have complex epistatic relationships with each other; for instance, the inactivation of a gene, complex, or pathway can make the inactivation of another gene in the same complex or pathway functionally redundant and thus neutral, and ii) the high number of mutations in a tumor can generate a large number of passenger mutations, even in driver genes^29, 51^.

To minimize these potential confounders, we first aligned the strength of causal effects in our mouse data with the frequency of alterations in human EGFR-driven lung adenocarcinoma. EGFR-driven lung adenocarcinomas have a low tumor mutational burden (TMB)^52^ (**Fig. 6a**), and our mouse data suggest that inactivation of several putative tumor suppressor genes are deleterious and thus unlikely to be observed in the human data, even as passengers. Indeed, there is a strikingly strong correlation between mouse cause-and-effect data and mutation frequencies in human EGFR-driven tumors (**Fig. 6b**; Spearman ρ=0.57, Pearson r=0.63). Restricting analysis of human mutation frequencies to patients with EGFR L858R mutation instead of patients with any EGFR-driven tumors in human data retains the strong correlation between mouse data and human mutational frequency data (Spearman ρ=0.57, Pearson r=0.67). The genes whose loss is predicted to be detrimental to EGFR-driven tumors (i.e., *Keap1, Lkb1*, and *Nf2*) are rarely co-mutated with EGFR in human lung adenocarcinoma as predicted (**Fig. 6b**). Furthermore, *P53* inactivation, which provides a strong benefit in our mouse models, is very commonly co-mutated with EGFR. Even excluding these extreme examples, the relationship between mouse causal data and human observational data remains strongly correlated (**Fig. 6b**; Spearman ρ=0.42, Pearson r=0.41).

**Figure 6.**
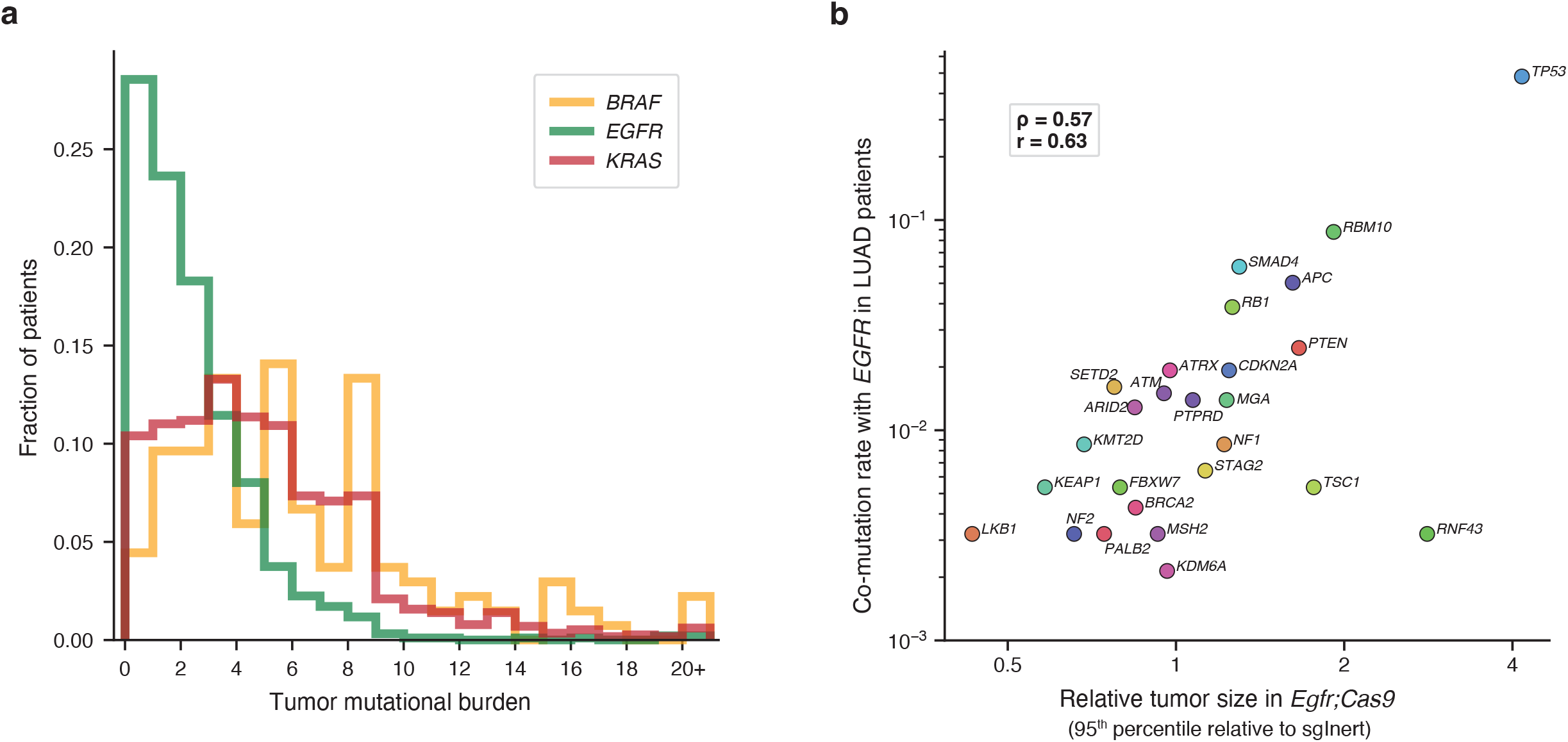
Causal effects predicted in mouse models correlate with frequency of alterations in EGFR-driven human lung adenocarcinoma, where most patients have low tumor mutational burden. **a.** Total number of mutations per patient per megabase in the LUAD cohort of the AACR Project GENIE database. Analysis was restricted to samples sequenced with the MSK-IMPACT468 panel. Missense, stop, and frameshift variants were included, and any mutations predicted by Polyphen as “benign” or by Sift as “tolerated” were excuded. Patient sample sizes were: *KRAS* N=1134, *EGFR* N=935, and *BRAF* N=135. The same patients were used for all following human analysis panels. **b.** Correlation of relative tumor size at the 95^th^ percentile to co-mutation rate of each gene tested in our model with *EGFR* in LUAD patients. *CMTR2* was the only gene tested in our model that was not present in the MSK-IMPACT468 panel and therefore not included in this analysis. Spearman rank-order correlation (ρ) and Pearson correlation (r) are indicated.

As anticipated, in the KRAS and BRAF contexts where the TMB is generally high there was a poor correlation between co-mutation frequency in human lung adenocarcinoma and causal mouse effects, suggesting that in these subgroups, human mutational frequency does not predict the importance of most tumor suppressor genes (**Supplementary Fig. 11a,b**). Consistent with previous studies, our analyses showed that in high-TMB tumors the mutation frequency of most tumor suppressor genes is strongly predicted by gene length and thus is very similar between the KRAS and BRAF contexts^53^ (**Supplementary Fig. 11c-d**, Spearman ρ=0.82, Pearson r=0.93). The implication of this observation is that most mutations, even those in functionally important tumor suppressor genes, in KRAS- and BRAF-mutant tumors are in fact passengers. High passenger mutation loads as well as a variety of mechanisms of tumor suppressor inactivation (beyond direct genomic alteration) together obscure functionally important interactions between oncogene and tumor suppressor alterations that are revealed by *in vivo* cause-and-effect experiments. For instance, *PTEN* is rarely mutated in KRAS- and BRAF-driven human lung cancers and yet *Pten* inactivation provides a very strong tumor fitness advantage in our autochthonous mouse models (**Fig. 6** and **Supplementary Fig. 11a,b**). Indeed, the PI3K pathway is commonly activated by non-mutational mechanism in human lung tumors, and *PTEN* and other members of the PI3K/AKT pathway are widely thought to be important regulators of human lung tumorigenesis^54^. This underscores the importance of unbiased functional genomic studies as we have done here, as driver alterations that occur rarely are not necessarily unimportant when present.

## DISCUSSION

In this study, we built an expansive fitness landscape of lung tumorigenesis by quantifying the joint effects of inactivating 28 known and putative tumor suppressor genes across four oncogenic contexts on tumor development *in vivo* (**Fig. 7**). In total, we quantified the fitness of 112 distinct oncogene by tumor suppressor pairs by assaying the ability of these genetic combinations to initiate tumorigenesis and drive tumor growth. While our previous work defined the fitness landscape of lung tumor suppression in the context of KRAS G12D^19^, the landscapes within the three other oncogenic contexts are largely novel (**Supplementary Fig. 1**). Going beyond understanding fitness within isolated oncogenic contexts, our multiplexed and quantitative approach allowed direct comparison of tumor suppressive effects across multiple contexts. Indeed, to our knowledge, we generated data on fifteen times more cross-oncogene tumor suppressor effect comparisons than have been studied previously in quantitative *in vivo* models (**Supplementary Fig. 12**). And although we did find alignment between growth effects in our model and human LUAD mutation rates in the context of *EGFR*—an oncogene notable for its low tumor mutational burden—most of the interactions we observed could not have been inferred from human data alone, e.g., with KRAS and BRAF mutation rates overwhelmed by passengers for about 80% of the genes studied.

**Figure 7.**
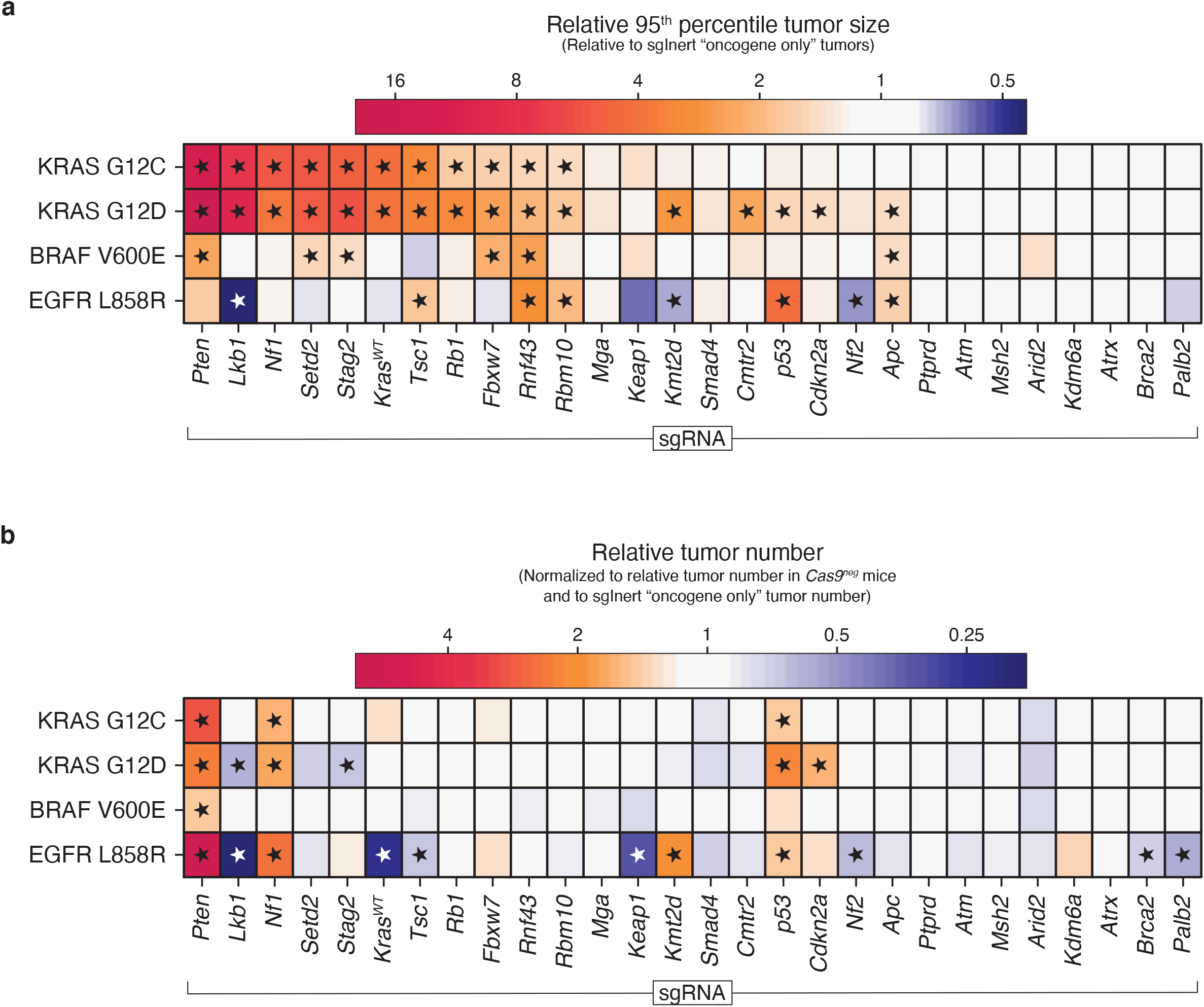
The impact of tumor suppressor pathways on tumorigenesis largely depends on which oncogene is activated, and is not predicted by the underlying strength of the oncogene alone. **a,b**. Relative tumor size ratio at the 95^th^ percentile (**a**) and relative tumor number (**b**) for tumors with the indicated Lenti-sgRNA/Cre vector on the x-axis and oncogenic allele on the y-axis. Asterisks indicate effects that are significant with FDR at 0.05 and also half of a log2-fold change from neutral.

The scale of our data allowed us to demonstrate that tumor suppressor effects vary strongly by oncogenic context and that the fitness landscape of tumor suppression displays strong and abundant epistasis. Of 28 tumor suppressors studied, few increased size or number across all oncogenes. Only inactivation of one, *Pten*, had a consistently strong, positive effect on both tumor growth and tumor number across all oncogenic contexts. *Rnf43* and *p53* consistently increased relative tumor size and number, respectively (**Fig. 7a,b**). Given the general trend of widespread epistasis, the robustness of the tumor suppression provided by these three genes is notable. The physiological role of *p53* in constraining tumor initiation/early expansion is striking and might be one reason for the prevalence and ubiquitous nature of *TP53* mutations in human cancer.

Many tumor suppressors showed clear sign epistasis with the oncogenes, whereby inactivation was advantageous in one context and either neutral or deleterious in another context. Surprisingly, inactivation of some of the strongest tumor suppressors in the presence of oncogenic KRAS variants decreased tumor growth in the presence of oncogenic EGFR. Furthermore, the oncogenic contexts were *qualitatively* different from each other: loss of tumor suppressors generally led to increased rates of tumor initiation and growth in the KRAS backgrounds, had more muted effects in the BRAF context, and had variable effects in the EGFR context.

Some of these epistatic effects were expected given our understanding of the RAS pathway and thus serve as positive controls. For example, the inactivation of NF1—a positive regulator of KRAS GTP to GDP transition—should shift KRAS proteins into their GTP-bound state and increase tumor number and/or growth in the KRAS G12C-, KRAS G12D- and EGFR-driven tumors. However, as class I mutations (e.g., BRAF V600E/D/K/R) have been demonstrated to activate MAPK signaling independent of upstream RAS signaling, inactivation of NF1 was expected to be neutral in BRAF V600E-driven lung tumors^55^. Likewise, inactivation of wild type KRAS was expected to increase tumor number and/or growth in the KRAS G12C and KRAS G12D contexts, as wild type KRAS suppresses oncogenic KRAS^34^. Conversely, inactivation of wild type KRAS was expected to reduce tumor number and/or growth in the EGFR context, as EGFR signals via wild type KRAS, and, as with NF1-deficiency, should not affect BRAF-driven tumors^21^. Indeed, we observed all of these expected effects in our data, providing an important validation of our results (**Fig. 2f,g, Fig. 4e, Fig. 7**, and **Supplementary Fig. 8d**).

The other cases of strong epistasis that we observed could not have been predicted based on the linear oncogenic EGFR➔KRAS➔BRAF pathway model. It is unclear why inactivation of *Lkb1, Setd2, Keap1, Kmt2d*, and *Nf2* leads to increased growth of oncogenic KRAS-driven lung tumors but is deleterious to EGFR-driven lung tumors (**Fig. 7**). This pattern indicates that the “off-axis” (i.e., not within the linear RAS pathway) signaling controlled by these three oncogenes drastically shifts the fitness effects of tumor suppressor losses. This should, in turn, affect the set of evolutionary trajectories that are likely after the initial oncogenic events of tumorigenesis.

Furthermore, the fitness effects of subsequent tumor suppressor inactivation could not have been predicted from the basal oncogenic potential of these four oncogenes, from strongest to weakest: KRAS G12D to BRAF V600E to KRAS G12C to EGFR L858R. The precise quantitative similarity of the tumor suppressive effects across KRAS variants is particularly striking because it implies that such effects can be robust to extreme (>10 fold) difference in oncogenic potential, differences in biochemical properties and enzymatic activities,^31, 35^ and diminishing-returns epistasis^7-9^. Overall, it appears that it is not possible to predict the impact of tumor suppressor alterations from simple linear pathway structures or from the fitness effects of the activated oncogenes in isolation.

This unpredictability may have implications for targeted therapeutic interventions, which work by repressing the signaling of oncogenes or co-linear nodes. If the epistasis also applies in reverse—i.e., that the fitness *costs* of oncogenic signal repression have strong, rugged interactions—then drug effects will be influenced by tumor suppressor inactivation in complex, target-specific ways that will require direct cause-and-effect empirical testing to unravel.

## ONLINE METHODS

### Design and generation of Lenti-sgRNA/Cre vectors

We generated lentiviral vectors encoding Cre (expressed from a PGK promoter^56^) and an sgRNA (expressed from a human U6 promoter) targeting each of the following 28 genes, which are known or putative tumor suppressors that are recurrently mutated in lung adenocarcinoma (or pan-carcinoma) and represent diverse cancer pathways^24, 36^: *Apc, Arid2, Atm, Atrx, Brca2, Cdkn2a, Cmtr2, Fbxw7, Kdm6a, Keap1, Kmt2d, Kras*^*WT*^, *Lkb1, Mga, Msh2, Nf1, Nf2, Palb2, Pten, Ptprd, Rb1, Rbm10, Rnf43, Setd2, Smad4, Stag2, Tsc1*, and *p53*. Vectors encoding “inert” sgRNAs were also generated: sg*Rosa26-1*, sg*Rosa26-2*, sg*Rosa26-3*, sg*NT-1*, sg*NT-2*, and sg*NT-3* were used in the *G12C;Cas9, G12D;Cas9*, and *Braf;Cas9* experiments, while sg*NT-2* and sg*Neo-1* were used in the *Egfr;Cas9* experiments.

sgRNAs were designed and selected as follows. Firstly, all possible 20-bp sgRNAs (using an NGG PAM) targeting each gene of interest were identified and scored for predicted on-target cutting efficiency using an available sgRNA design/scoring algorithm^57^. For each tumor suppressor gene, we then selected the sgRNA predicted to be the most likely to produce null alleles: preference was given to sgRNAs that were previously validated *in vivo*^29, 31, 58^, had the highest predicted on-target cutting efficiencies, targeted exons conserved in all known splice isoforms (ENSEMBL), targeted splice acceptor/splice donor sites, positioned earliest in the gene coding region, occurring upstream of or within annotated functional domains (InterPro; UniProt), and occurring upstream of or at known recurrent mutation sites in human lung adenocarcinomas. The sgRNA sequences for each target are listed below:

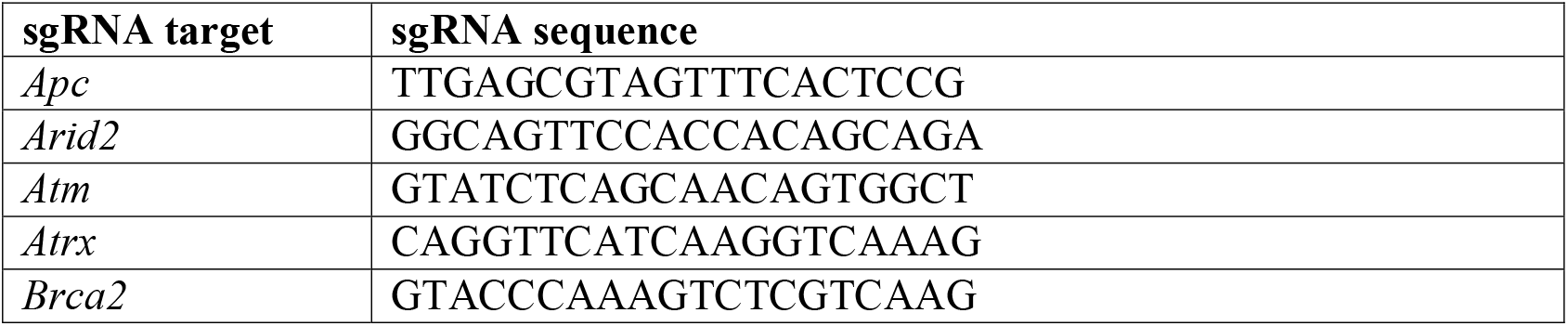
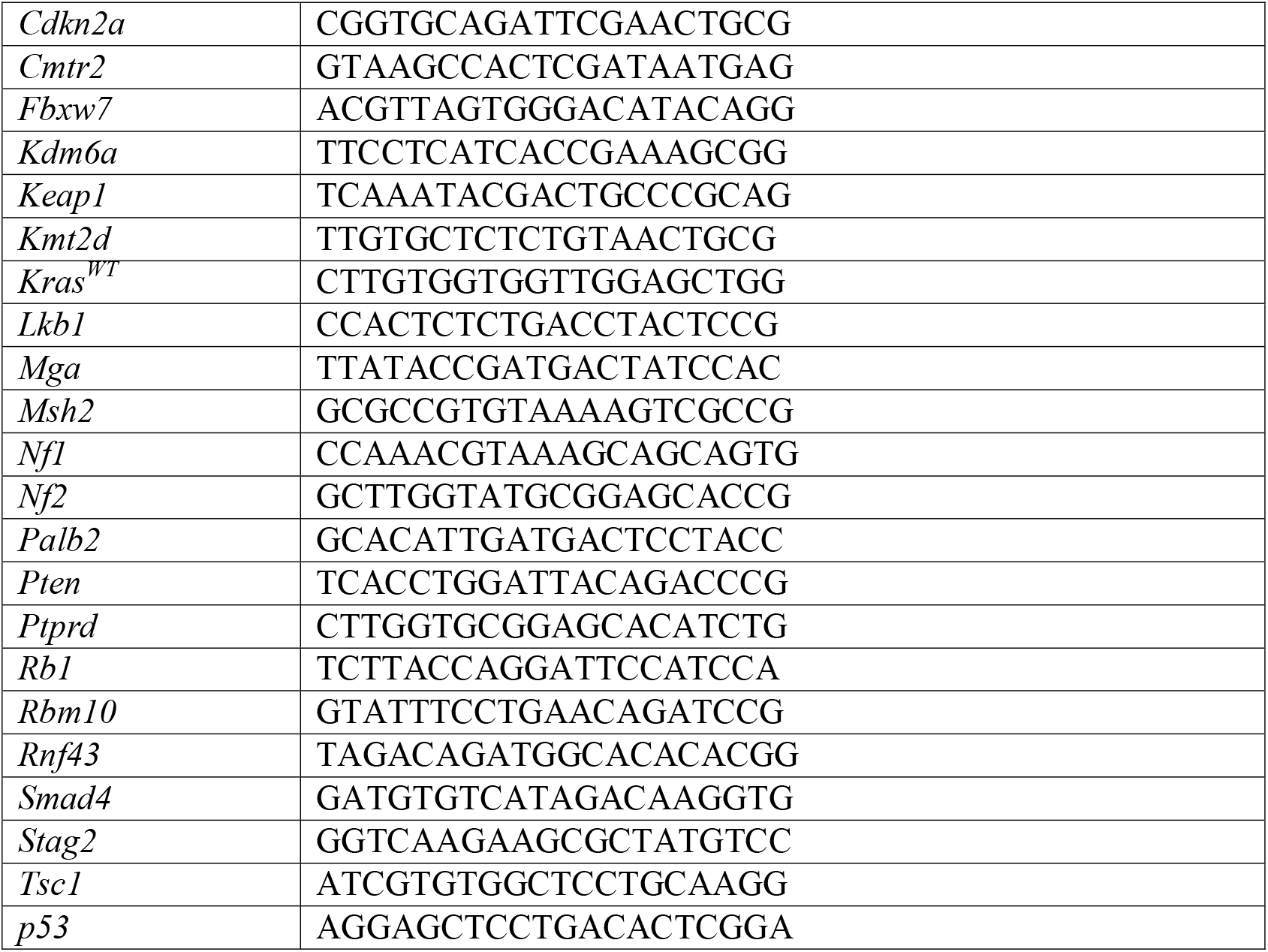

To generate Lenti-sgRNA/Cre vectors containing each sgRNA, a double-stranded DNA fragment (IDT gBlock) containing a U6-sgRNA-tracrRNA cassette flanked by restriction sites (AscI and SbfI) was synthesized and digested by AscI and SbfI. This digested DNA fragment was then cloned into an AscI/SbfI-digested parental lentivector encoding Cre to produce each circularized Lenti-sgRNA/Cre vector.

### Barcode diversification of Lenti-sgRNA/Cre

To enable quantification of the number of cancer cells in individual tumors in parallel using high-throughput sequencing, we diversified the Lenti-sgRNA/Cre vectors with a 46 bp multi-component barcode cassette that would be unique to each tumor by virtue of stable integration of the lentiviral vector into the initial transduced cell. This 46 bp DNA barcode cassette was comprised of a known 6-nucleotide ID specific to the vector backbone (vectorID), a 10-nucleotide ID specific to each individual sgRNA (sgID), and a 30-nucleotide random barcode containing 20 degenerate bases (random BC) (**Supplementary Fig. 2a**).

The 46 bp barcode cassette for each sgRNA was flanked by universal Illumina TruSeq adapter sequences and synthesized as single-stranded DNA oligos. Forward and reverse primers complimentary to the universal TruSeq sequences and containing 5’ tails with restriction enzyme sites (AscI and NotI) were used in a PCR reaction to generate and amplify double-stranded barcode cassettes for cloning. Each Lenti-sgRNA/Cre vector and its matching insert barcode PCR product was digested with AscI and NotI.

To generate a large number of uniquely barcoded vectors, we ligated 1 µg of linearize vector and 50 ng of insert with T4 DNA ligase in a 100 µl ligation reaction. 4-5 hours after incubation at room temperature, ligated DNA was precipitated by centrifugation at 14,000 rpms for 12 minutes after adding 5 µl Glycogen (5 mg/ml) and 280 µl 100% Ethanol into the ligation reaction. The DNA pellet was washed with 80% Ethanol and air-dried before being resuspended with 10 µl water. This 10 µl well-dissolved DNA was transformed into 100 µl of Sure Electrical Competent Cells using BioRad electroporation system following manufacturer’s instructions. Electroporation-transformed cells were immediately recovered by adding into 5 ml pre-warmed SOC media. From these 5 ml of bacteria, 10 µl were further diluted with LB ampicillin broth, and a final dilution of 1:200,000 was plated on an LB ampicillin plate for incubation at 37ºC. The remaining bacteria were mixed gently and thoroughly before being inoculated into 100 ml LB/Ampicillin broth, shaking at 220 rpm at 37ºC overnight. The next day, colony numbers on the LB/Ampicillin plate were counted to estimate the complexity of each library and the 100 ml bacterial culture was pelleted for plasmid purification.

8 colonies from each library were picked and PCR screened for verification of the specific sgRNA sequence and corresponding barcode sequence among these 8 colonies. The final purified library plasmid for each library was again sequence verified.

### Production, purification, and titering of lentivirus

24 hours prior to transfection, 2.4×10^7^ 293T cells were plated on a 15 cm tissue culture plate. 30 µg of pPack (packaging plasmid mix) and 15 µg of library plasmid DNA were mixed well in 1.5 ml serum free D-MEM medium before an equal volume of serum free D-MEM medium containing 90 µl of LipoD293 was added. The resulted mixture was incubated at room temperature for 10-20 minutes before adding into 293T cells. At 24 hours post-transfection, replace the medium containing complexes with 30 ml of fresh D-MEM medium supplemented with 10% FBS, DNase I (1 unit/ml), MgCl2 (5 mM), and 20 mM HEPES, pH 7.4. The entire virus-containing medium from each plate was collected and filtered through a 0.2 µm PES filter (Nalgene) at 48 hours post-transfection. The viruses were further concentrated by centrifugation at 18,500 rpm, 4ºC for 2 hours, and the pellet was resuspended in 500 µl PBS buffer. 50 µl virus aliquots were stored at -80°C.

To quantify the titer of packaged library constructs, 10^5^ LSL-YFP MEF cells^28^ were transduced with 1 µl of viruses in 1 ml culture medium containing 5 µg/ml polybrene. Transduced cells were incubated for 72 hours before being collected for FACS analysis to measure the percentage of YFP-positive cells. Control viruses were used in parallel to normalize the virus titers.

### Pooling of Lenti-sgRNA/Cre vectors

To generate a pool of barcoded Lenti-sgRNA/Cre vectors for initiation of multiple tumor genotypes within individual mice, barcoded Lenti-sgRNA/Cre vectors targeting the 28 genes described above (*Apc, Arid2, Atm, Atrx, Brca2, Cdkn2a, Cmtr2, Fbxw7, Kdm6a, Keap1, Kmt2d, Kras*^*WT*^, *Lkb1, Mga, Msh2, Nf1, Nf2, Palb2, Pten, Ptprd, Rb1, Rbm10, Rnf43, Setd2, Smad4, Stag2, Tsc1*, and *p53*), and those containing the inert, negative control sgRNAs, were combined such that the viruses would be at equal ratios in relation to their estimated in vitro or in vivo titers. In the *EGFR;Cas9* experiment, some viruses were underrepresented in the pool as we were limited by total volume of those viruses, and in that same experiment, the virus pool contained additional targets for which data were not included in this study. All virus pools were diluted with 1x DPBS to reach the necessary titer for each experiment.

### Mice, tumor initiation, and tissue collection

*Kras*^*LSL-G12D*^, *Braf*^*CA-V600E*^, *tetO-EGFR*^*L858R*^, *Rosa26*^*LSL-rtTA3-ires-mKate*^, *Rosa26*^*LSL-Cas9-2a-GFP*^ and *H11*^*LSL-Cas9*^ alleles have been described^28, 38, 39, 59-61^. Lung tumors in all mice were initiated via intratracheal delivery of a lentivirus pool containing barcoded Lenti-sgRNA/Cre vectors targeting 28 genes (*Apc, Arid2, Atm, Atrx, Brca2, Cdkn2a, Cmtr2, Fbxw7, Kdm6a, Keap1, Kmt2d, Kras*^*WT*^, *Lkb1, Mga, Msh2, Nf1, Nf2, Palb2, Pten, Ptprd, Rb1, Rbm10, Rnf43, Setd2, Smad4, Stag2, Tsc1*, and *p53*) plus 6 negative control sgRNAs in the *G12D;Cas9, G12C;Cas9*, and *Braf;Cas9* experiments (three targeting the *Rosa26* gene, which are actively cutting but functionally inert, and 3 non-cutting sgRNAs with no expected genomic target [sgNon-Targeting: sg*NT*]) and 2 negative control sgRNAs in the *Egfr;Cas9* mice (one targeting the Neomycin resistance gene within the *Rosa26* allele, which is actively cutting but functionally inert^61^, and one non-cutting sgRNA with no expected genomic target [sgNon-Targeting: sg*NT*]). To induce oncogenic EGFR expression in *Egfr;Cas9* mice, mice were fed doxycycline-impregnated food pellets (625 ppm; HarlanTeklad) starting 1-2 days prior to delivery of pooled barcoded Lenti-sgRNA/Cre vectors.

Whole lung tissue was extracted from euthanized mice as previously described^58^. Lung mass measurements were recorded as a proxy for overall lung tumor burden. Individual lung lobes from some mice were inflated with 10% neutral buffered formalin and allowed to fix for 16-24 hours before passaging into 70% Ethanol for subsequent embedding, sectioning, and histological analyses using conventional methods. Remaining lung tissue was weighed and then stored at -80ºC prior to subsequent processing for next-generation sequencing (see sections below).

All animals were kept in pathogen-free housing and animal experiments were conducted in accordance with protocols approved by either the Yale University Institutional Animal Care or Explora BioSciences Institutional Animal Care and Use Committee (IACUC) guidelines.

### Generation of spike-in controls

DNA barcode cassettes comprised of 46 bp barcode cassettes and flanked by universal Illumina TruSeq adapter sequences as well as additional buffer sequences to extend their total length to >400bp were generated either by direct synthesis of the double-stranded DNA fragments (GeneWiz, IDT) or synthesis of single-stranded DNA oligos (GeneWiz, IDT) with overlapping complementary regions that were extended and amplified via PCR to create double-stranded DNA products that were then purified. Aliquots of these stock double-stranded DNA fragments were diluted to the desired copy numbers using DNase-free ultra-pure H2O and stored at -20ºC.

### Isolation of genomic DNA from mouse lungs

Whole lungs were removed from freezer and allowed to thaw at room temperature. Spike-ins were added to each whole lung samples. Qiagen Cell Lysis Buffer and proteinase K from Qiagen Gentra PureGene Tissue kit (Cat # 158689) was added as described in the manufacturer protocol. Whole lungs plus spike-ins from each mouse were homogenized in the Cell Lysis buffer and Proteinase K solution using a tissue homogenizer (FastPrep-24 5G, MP Biomedicals Cat # 116005500). Homogenized tissue was incubated at 55ºC overnight. To remove RNA from each tissue samples, RNase A was added with additional spike-ins to whole homogenized tissue. To maintain an accurate representation of all tumors, DNA was extracted, and alcohol precipitated from the entire lung lysate using the Qiagen Gentra PureGene kit as described in manufacturer protocol. More spike-ins were added to the resuspended DNA.

### Preparation of barcode libraries for sequencing

Libraries were prepared by amplifying the barcode region from 32 µg of genomic DNA per mouse. The barcode region of the integrated Lenti-sgRNA/Cre vectors was PCR amplified using primer pairs that bound the universal Illumina TruSeq adapters and contained dual unique multiplexing tags. We used a single-step PCR amplification of barcode regions, which we found to be a highly reproducible and quantitative method to determine the number of cancer cells in each tumor. We performed eight 100 µl PCR reactions per mouse (4 µg DNA per reaction) using Q5 HF HS 2x mastermix (NEB #M0515) with the following PCR program:

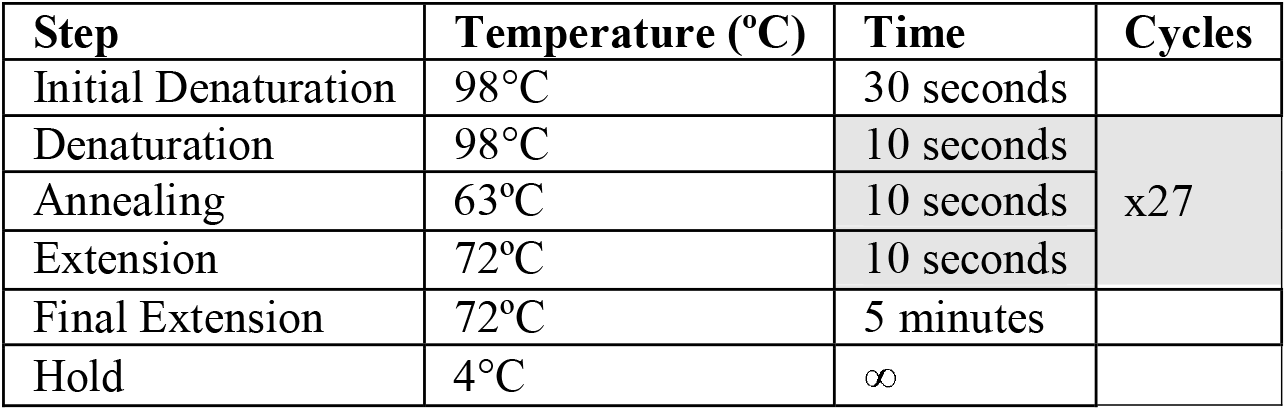

| Step | Temperature (°C) | Time | Cycles |
| --- | --- | --- | --- |
| Initial Denaturation | 98°C | 30 seconds |  |
| Denaturation | 98°C | 10 seconds | x27 |
| Annealing | 63°C | 10 seconds |  |
| Extension | 72°C | 10 seconds |  |
| Final Extension | 72°C | 5 minutes |  |
| Hold | 4°C | ∞ |  |

The concentration of amplified barcode product in each PCR was determined by TapeStation (Agilent Technologies). Sets of 20-60 PCR were pooled at equal molar ratios of barcode product, normalized to the estimated burden of tumors in each mouse lung sample (measured lung mass minus an estimated normal lung mass of between 0.15 and 0.18g) associated with the PCRs. Pooled PCRs were cleaned up using a two-sided SPRI bead purification. Samples were sequenced on an Illumina NextSeq.

### Analysis of sequencing data

Paired-end sequencing reads were demultiplexed via unique dual indexes using BCLConvert (version 3.8.2) and adapters sequences were trimmed using CutAdapt (version 4.1). CutAdapt was used in paired-end mode with the following parameters: minimum-length=0, error-rate=0.1, overlap=3. Paired-end alignments were constructed between mate-paired reads and library-specific databases of the expected oligonucleotide spike-in and tumor barcode insert sequences using Bowtie2 (version 2.4.4). These alignments were stringently filtered from downstream analysis if they failed to meet any of several quality criteria, including:

- No mismatches between the two mate-pairs, which fully overlap one another, at any location.
- No mismatches between the mate-paired reads and expected constant regions of the barcode or spike-in to which they best align.
- No indels in alignments between mate-paired reads and the barcode or spike-in to which they best align.

Following alignment, errors in paired-end reads were corrected via a simple greedy clustering algorithm:

- Reads were *dereplicated* into read sequence/count tuples, (s_i_, r_i_)
- These tuples were re-ordered from highest to lowest based on their read abundances, {r_i_}.
- This list of tuples was traversed from i = 1…N, taking one of the following actions for each tuple (s_i_, r_i_):
  ∘ If s_i_ *is not* within a Hamming distance of 1 from any s_j_ with j < i, then (s_i_, r_i_) initiates a new cluster.
  ∘ If s_j_ *is* within a Hamming distance of 1 from some s_j_ with j < i, then it joins the cluster of s_j_.

The resulting clusters are each considered to represent an error-corrected sequence equal to that of the sequence that founded the cluster with read count equal to the sum of the read counts of the dereplicated reads that are members of the cluster.

A second stage of error correction was performed to remove additional errors. Hamming distance D(s_i_, s_j_) was computed on all pairs of error-corrected sequences. Then, each sequence s_i_ (with r_i_ reads) was absorbed into the most abundant sequence s_j_ (with r_j_ > r_i_ reads) if either of the following criteria were met:

- D(s_i_, s_j_) ≤ 3
- D(s_i_, s_j_) ≤ 5 and r_j_/r_i_ ≥ 5 or r_i_ ≤ 3

These heuristics were established based on internal control data. After applying both rounds of error correction, we estimate a false positive rate of 1.4×10^−8^ based on the number of reads assigned to spike-in oligonucleotide sequences (which have no degenerate bases) that were not added to the samples. Following error correction, a filter was applied to remove sequences that could have originated from cross-contamination: barcodes were compared across samples in the same study, and any exact sequences that were found in more than one library were removed.

Following error correction and cross-contamination removal, the read counts of each unique barcode were converted to neoplastic cell number by dividing the number of reads of the spike-in oligonucleotide added to the sample prior to tissue homogenization and lysis at a fixed, known concentration.

### Removal of mice that did not get sufficient viral titer during transduction

Following the sequence processing, mice were removed if they did not reach a lower bound of total neoplastic cells. For the experiments with *G12D;Cas9* and *G12C;Cas9*, mice were removed if they had less than 10^6^ total neoplastic cells. For the experiments with *Braf;Cas9* and *Egfr;Cas9* mice, mice were removed if they had less than 10^5^ total neoplastic cells. Thresholds were chosen using by examining the distribution of total neoplastic cells per mouse across each study. Most mice fall within ∼2 orders of magnitude of each other, and any outliers fell at least an order of magnitude below the rest of the distribution.

### Accounting for processed lung mass when normalizing metrics by titer

Because several mice had lobes taken for histology and therefore only a fraction of the lung made it into TubaSeq, the processed lung should not be expected to represent the full viral titer transduced to the mouse. To correct the titer for that fraction of lung, we multiplied the total titer given to the mouse by the ratio of the processed lung weight to the total lung weight before any lobes were removed. This effective titer was used for all plots that present titer-normalized quantities (e.g., **Fig. 1f-i, 3d-g**).

### Calculation of tumor size percentiles

First, tumors were pooled across all mice in the group, and separated into tumors that map to each Lenti-sgRNA/Cre guide. Tumors from Lenti-sgInert/Cre were pooled (sg*NT*-1, sg*NT*-2, sg*NT*-3, sg*R26*-1, sg*R26*-2, sg*R26*-3 for experiments using *G12D;Cas9, G12C;Cas9, Braf;Cas9* mice, and sg*NT*-2, sg*Neo*-1 for the experiments using *Egfr;Cas9* mice) to create one pool of sgInert tumors. Using the sgInert tumors, a minimum tumor size cutoff was determined, above which tumor percentiles would be calculated. The goal of matching this cutoff across study groups, particularly when comparing across oncogenes, was to ensure that the tumor suppressor effects were being measured on the same fraction of initiated tumors, independent of the strength of the oncogene. A cutoff was chosen for each study group that matched the number of sgInert tumors per titer above the cutoff. The exception was for the *Braf;Cas9* experiments. Because the *Braf;Cas9* tumor sizes differed so strikingly from those in the *G12D;Cas9, G12C;Cas9, Egfr;Cas9* experiments, suggesting a very different process for tumor initiation and/or growth, we opted to use an *ad hoc* cutoff that captured >85% of total tumor burden and reduced the high mouse-to-mouse variability in the number of small tumors. For the main experiments where all mice of each oncogene-timepoint pair were pooled, the following minimum cutoffs were used: 1600 cells for *G12D;Cas9* 15 weeks, 600 cells for *G12D;Cas9* at 9 weeks, 400 cells for *G12C;Cas9* at 15 weeks, 300 cells for *G12C;Cas9* at 9 weeks, 300 cells for *Egfr;Cas9* at 15 weeks, and 3000 cells for *Braf;Cas9* at 15 weeks. Neoplastic cell number cutoffs for the comparisons of the replicate study groups in **Fig. 2** were calculated using the same procedure.

For each set of Lenti-sgRNA/Cre tumors in each oncogene-background pair, size percentiles of tumors above the cutoff were computed and divided by the same size percentiles for the sgInert tumors in the same context with the same cutoff. This ratio is referred to as *relative tumor size* in **Fig. 2, 4, and 6** and **Supplementary Fig. 4, 6, and 8**. Mice and tumors were bootstrapped 8000 times and calculation was repeated each time. A 95% confidence interval from these bootstraps was reported.

### Calculation of tumor number enrichment

Tumor number enrichment estimates the factor by which there are more or fewer sgRNA tumors above a minimum size cutoff than there would have been if the tumors had been sgInert. As we sought to measure tumor number effects associated with initiation and very early tumor growth, a cutoff of 500 cells was used, as it represents a lower bound on our technical resolution. First, all tumors from each mouse in each study group being compared are pooled and separated into tumors that map to each Lenti-sgRNA/Cre guide. Tumors from Lenti-sgInert/Cre were pooled (sg*NT-*1, sg*NT*-2, sg*NT*-3, sg*R26*-1, sg*R26*-2, sg*R26*-3 for experiments using *G12D;Cas9, G12C;Cas9, Braf;Cas9* mice, and sg*NT*-2, sg*Neo*-1 for the experiments using *Egfr;Cas9* mice) to create one pool of sgInert tumors. Because tumor number for a given sgRNA will be proportional to the titer of the individual sgRNAs within the viral pool, we calculated the ratio of sgRNA tumors to sgInert tumors in mice without Cas9 in each virus pool, which is expected to be driven only by titer differences between viruses.

Then, to calculate the tumor number enrichment for each sgRNA, we divided the number of sgRNA to sgInert tumors, and divided this ratio by the same ratio in the mice without Cas9 that had been initiated with the same pool of viruses. Log_2_ of this ratio of ratios is referred to as *relative tumor number* in **Fig. 5 and 6** and **Supplementary Fig. 9 and 10**. Mice and tumors were bootstrapped 8000 times and calculation was repeated each time. A 95% confidence interval from these bootstraps was reported.

### Calculation of tumor burden densities

The density of tumor burden as a function of log tumor size was estimated as follows. First, we pooled tumors across all mice in each cohort and computed total tumor burden by summing the sizes of all tumors of all sizes. We then generated log-spaced bins with 10 bins per order of magnitude of tumor size, summed the sizes of all Lenti-sgInert/Cre tumors in each bin, and divided by total tumor burden. To create a density in log size, we then divided this ratio by log (bin width), which was a constant given the log-spaced binning. Finally, mice and tumors were bootstrapped 1000 times and this procedure repeated each time, and the mean density across bootstraps as well as a 95% confidence interval are shown for each size.

### Calculating lengths of gene coding regions

For each gene of interest, we used the coding sequence annotations of the “Ensembl Canonical” transcript (Ensembl project, release 105)^62^ to determine the length of the gene’s coding region.

### Determining gene mutation and co-mutation rates from human lung cancer genomics data

Mutation rates in human LUAD were estimated using AACR Project Genie (release version Genie 12.0) ^25^. First, we restricted our analysis to patients with LUAD and then selected those with the relevant oncogene mutations. We used the following definitions of *KRAS, EGFR*, and *BRAF* oncogene mutations:

- *KRAS*: any mutation in codon 12, 13, or 61
- *EGFR*: p.L858R, p.L861Q, p.G719X, deletion or insertion in exon 19, insertion in exon 20
- *BRAF*: all mutations listed in Table 1 of Owsley et al.^63^

To minimize bias due to the variety of genetic panels used in AACR Project Genie, we restricted our analysis to patients sequenced with the MSK-IMPACT468 panel, which was the most commonly used panel for LUAD patients. This resulted in 1134 patients with *KRAS* mutations, 935 with *EGFR* mutations, and 135 with *BRAF* mutations. *CMTR2* was the only gene that we tested that was not included in the panel, so we excluded it from our analysis. Correlations of mutation frequencies and mouse effects were also assessed using all panels in the AACR Project Genie database, and Spearman and Pearson correlations produced were similar to those using only MSK-IMPACT468.

To determine the co-mutation frequencies for each of the tumor suppressor genes we inactivated in our mouse models, we counted mutations as follows: first, we selected all mutations that were nonsense, missense, or frameshift variants. Then, any mutations that were predicted by Polyphen to be benign or predicted by SIFT to be tolerated were excluded. Correlations between causal mouse effects and human co-mutation frequencies were also tested with other definitions, and we confirmed that as we move from the set of mutations defined above to non-synonymous mutations, and then to all mutations the Spearman correlations decreased slightly, but the trend remained the same.

Tumor mutational burden was calculated as the total number of mutations per 1,000,000 base pairs. The total gene length was calculated using the sum of exon length for all genes that were queried in the panel.

When correlating exon length and co-mutation frequency, outliers were first removed. We found the same set of outliers using two methods: 1) Clustering was run using the pam algorithm, a robust version of Kmeans, and the gap statistic was calculated using the “globalSEmax” method, which looks for the first value that is lower than the global maximum minus the standard error when evaluated at that value. 2) Spearman correlation was calculated using the full set of genes, and then genes were removed one at a time to test how the correlation increased. Each iteration, the gene whose removal maximized the Spearman correlation was removed. A plot of number of genes removed against Spearman correlation showed an elbow at the removal of 5 genes. Both methods determined *CDKN2A, TP53, KEAP1, LKB1*, and *RBM10* to be the outliers. Linear regression on a log-log scale was then performed on the remaining genes. Pearson correlation and Spearman correlation were reported on the plot.

Relative tumor size at the 95^th^ percentile was used as the metric of mouse causal effects. Correlation of co-mutation frequency in *EGFR*-driven tumors with relative tumor size at 90^th^, 75^th^, and 50^th^ percentiles maintained the trend but became less strong as the percentiles increased, with Spearman correlations of 0.53, 0.49, and 0.43, respectively.

## ACKNOWLEDGEMENTS

We thank Explora and The Jackson Laboratory for expert animal care. We are grateful to all members of D2G Oncology. for expert advice and helpful comments. We would like to acknowledge the American Association for Cancer Research and its financial and material support in the development of the AACR Project GENIE registry, as well as members of the consortium for their commitment to data sharing (interpretations are the responsibility of study authors). We thank AstraZeneca, Bristol-Myers Squibb, Revolution Medicines, Merck KGaA, and CureTeq for generously allowing the inclusion of data generated in collaboration with D2G Oncology. L.E.D. is the Burt Gwirtzman Research Scholar in Lung Cancer. D.A.P. and M.M.W. are supported by NIH R01-CA 234349. This work was supported in part by NIH SBIR R44-CA250672.

---

**Supplementary Table 1.**
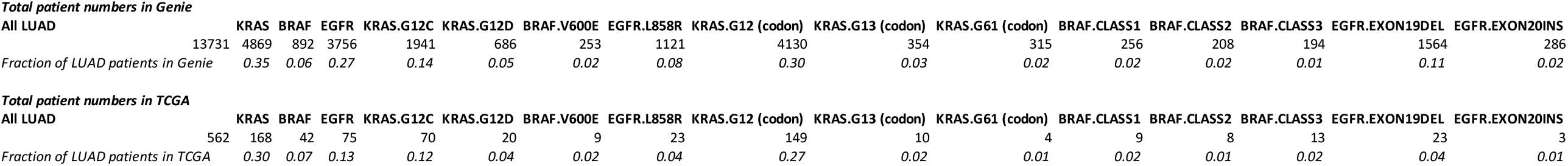
Frequency of oncogenic KRAS, EGFR, and BRAF mutations in LUAD.

**Supplementary Figure 1.**
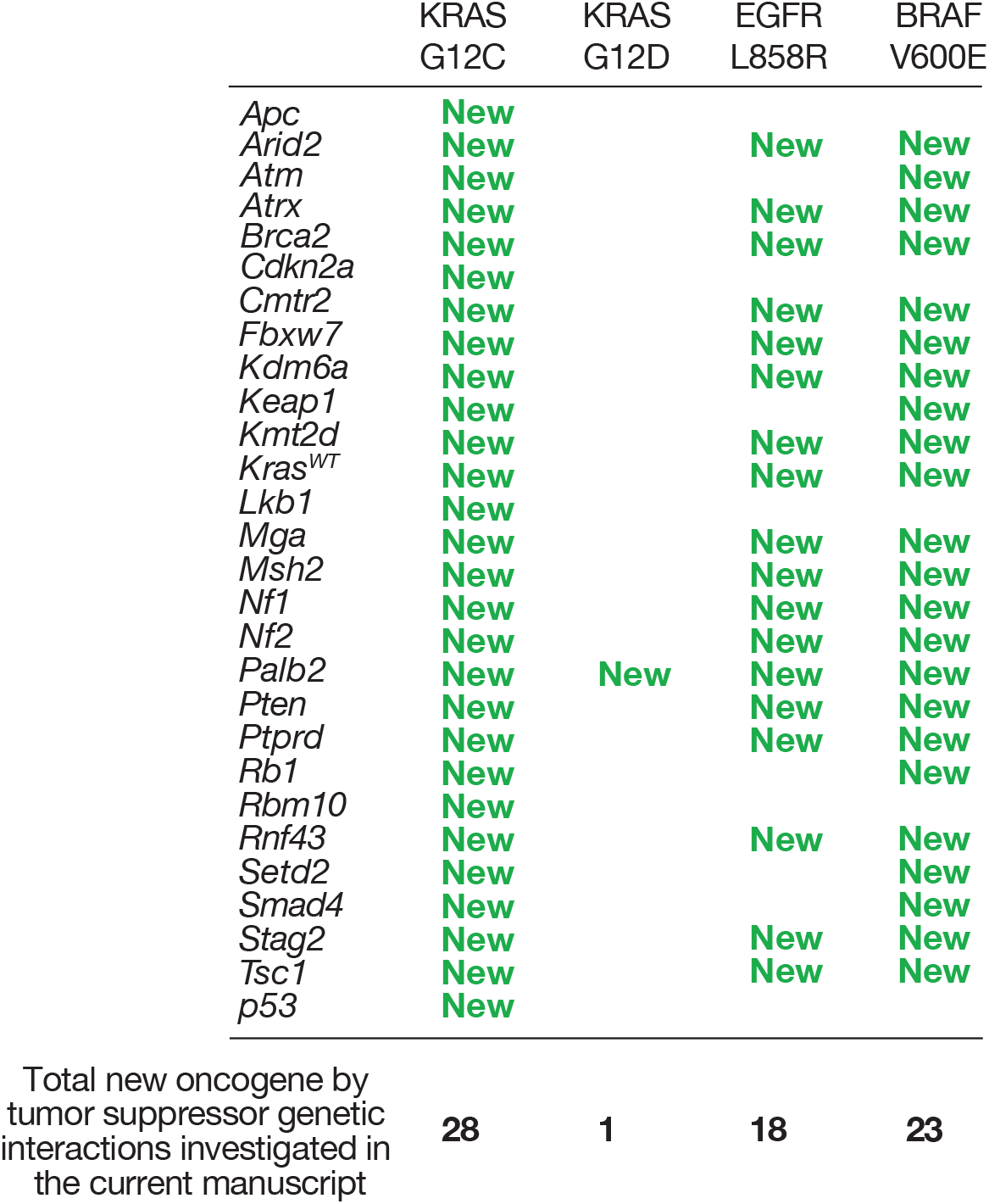
Extent of novel oncogene by tumor suppressor genetic interactions described in the current manuscript. Previous studies have investigated the impact of inactivating putative tumor suppressor genes on lung tumor growth in similar genetically engineered models. “New” indicates that this oncogene by tumor suppressor combination has not been previously investigated. The number of new oncogene by tumor suppressor combinations analyzed in this manuscript is indicated at the bottom of each column (70 out of the 112 possible combination have not been investigated previously). Many of the genes investigated in the oncogenic KRAS G12D-driven model have been investigated using conventional tumor suppressor floxed alleles as well as using Tuba-seq (Roger *et al*., 2017 and Cai *et al*., 2021). All of the tumor suppressors analyzed in the model of EGFR-driven lung cancer were performed using Tuba-seq (Foggetti *et al*., 2021). The tumor suppressors studied in the model of BRAF-driven lung cancer have all been analyzed using conventional floxed models, hence there has been no systematic cross comparison of tumor suppressor effects within that model. See **Supplementary Fig. 12** for a summary of existing data that compared tumor suppressor effects across (rather than within) genetically engineered lung cancer mouse models.

**Supplementary Figure 2.**
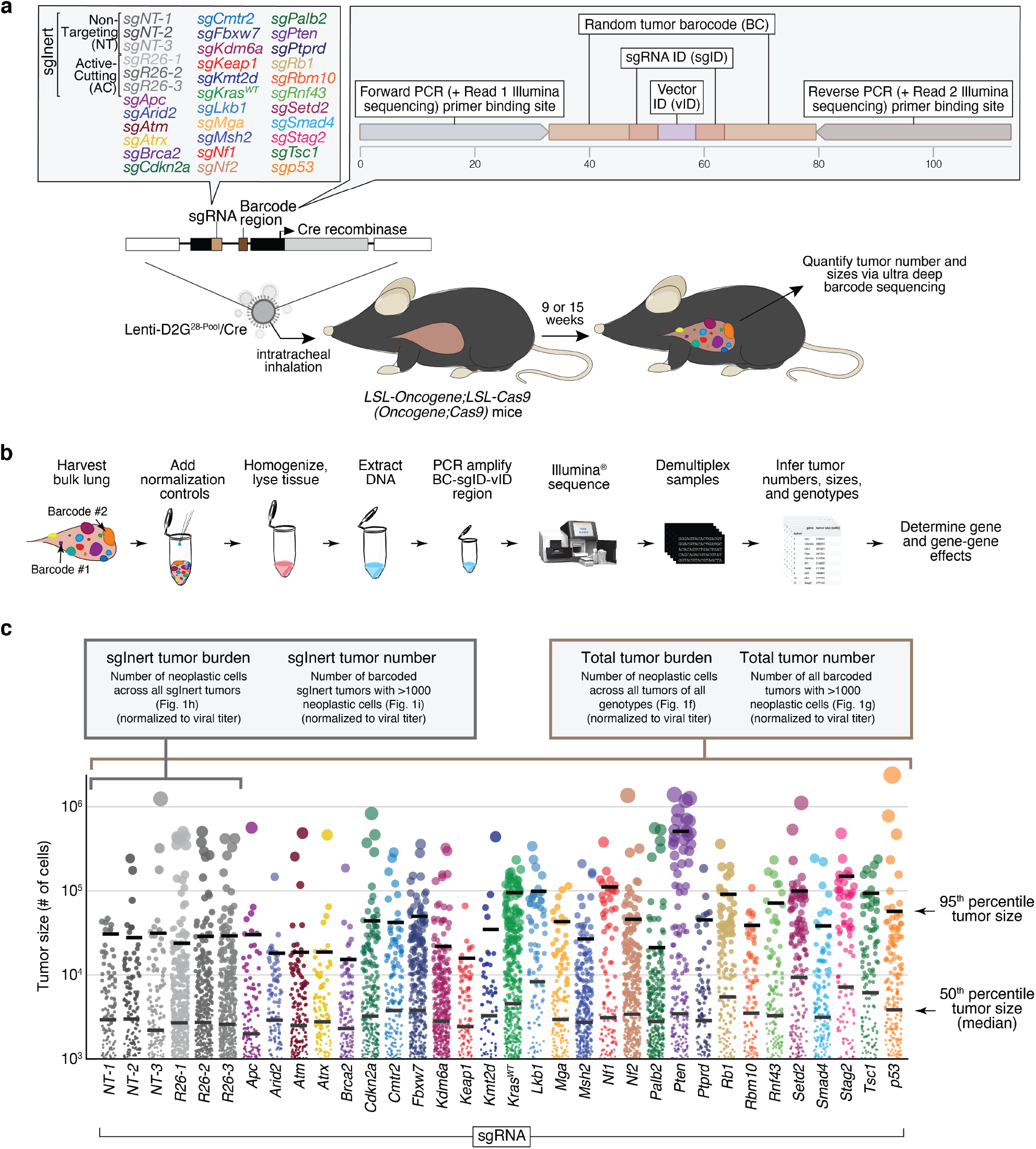
Integration of CRISPR/Cas9-mediated gene editing and tumor barcoding into genetically engineered mouse models enables the generation of tumors with diverse combinations of activated oncogenes and inactivated tumor suppresors within individual mice. **a.** General experimental schematic depicting the composition of the pool of barcoded Lenti-sgRNA/Cre vectors (Lenti-D2G^28-Pool^/Cre) and composition of the three component barcode. **b.** Overview of sample processing starting from bulk lung. The addition of barcoded normalization controls at a known number allows the number of neoplastic cells in each tumor to be calculated (see Methods). **c.** Jitter plot of all barcoded tumors detected in one *G12D;Cas9* mouse with Lenti-D2G^28-Pool^/Cre initiated tumors. Each dot represents a tumor and the dot size is scaled to tumor size. The size of the 95^th^ percentile and 50^th^ percentile (median) tumor within the size distribution of tumors with each Lenti-sgRNA/Cre vector is indicated. Metrics of total tumor burden, total tumor number, sgInert tumor number, and sgInert tumor burden are shown.

**Supplementary Figure 3.**
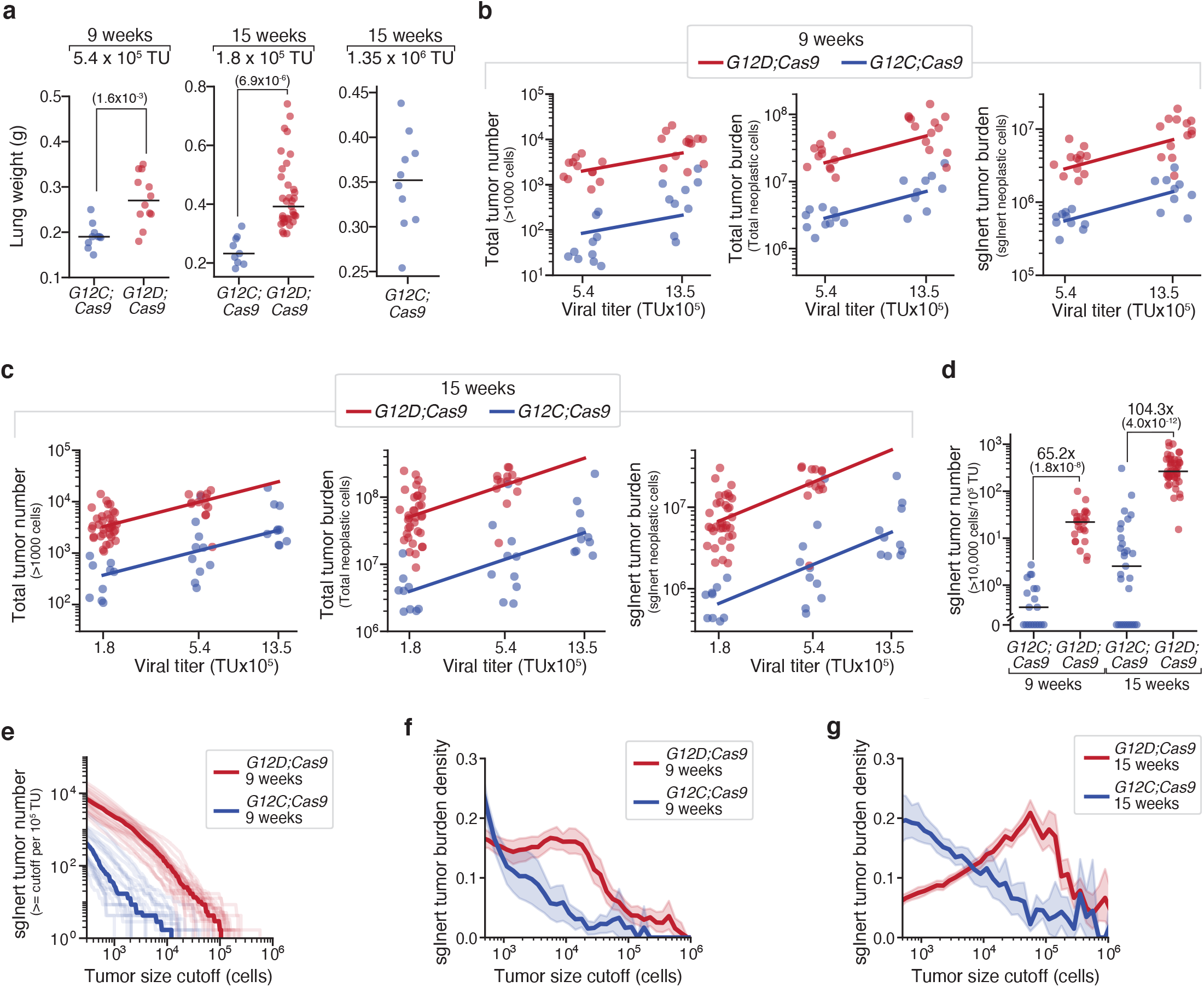
KRAS G12C consistently generates lower tumor burden than KRAS G12D. **a**. Lung weights of mice transduced with the indicated titers of Lenti-D2G^28-Pool^/Cre. Genotype and time post-tumor initiation is indicated. Each dot represents a mouse and the bar is the median. Fold difference between medians and significance calculated using a wilcoxon rank sum test (p-values < number in parentheses) is shown. **b**. Total tumor number, total tumor burden, and sgInert tumor burden number of neoplastic cells in mice 9 weeks post-tumor initiation with the indicated titer of virus. Mouse genotypes are indicated. Each dot represents a mouse and the bar is the median. Lines represent a linear fit through the origin, showing the increases in tumor number and burden are linear with titer. **c**. Total tumor number, total tumor burden, and sgInert tumor burden number of neoplastic cells in mice 15 weeks post-tumor initiation with the indicated titer of virus. Mouse genotypes are indicated. Each dot represents a mouse and the bar is the median. Lines represent a linear fit through the origin, showing the increases in tumor number and burden are linear with titer. **d**. Number of large sgInert tumors greater than 10,000 cells in size, normalized to viral titer. Mouse genotypes and time-points are indicated. Each dot represents a mouse and the bar is the median. Fold difference and significance calculated using a wilcoxon rank sum test (p-values < number in brackets) are shown. **e**. Number of tumors at or above the tumor size cutoff on the x-axis in *G12D;Cas9* and *G12C;Cas9* at 9 weeks post-tumor initiation. Each transparent line represents a mouse and the solid line is the median. **f,g**. The density function of sgInert tumor burden as a function of log(tumor size) 9 weeks (**f**) and 15 weeks (**g**) post-tumor initiation.

**Supplementary Figure 4.**
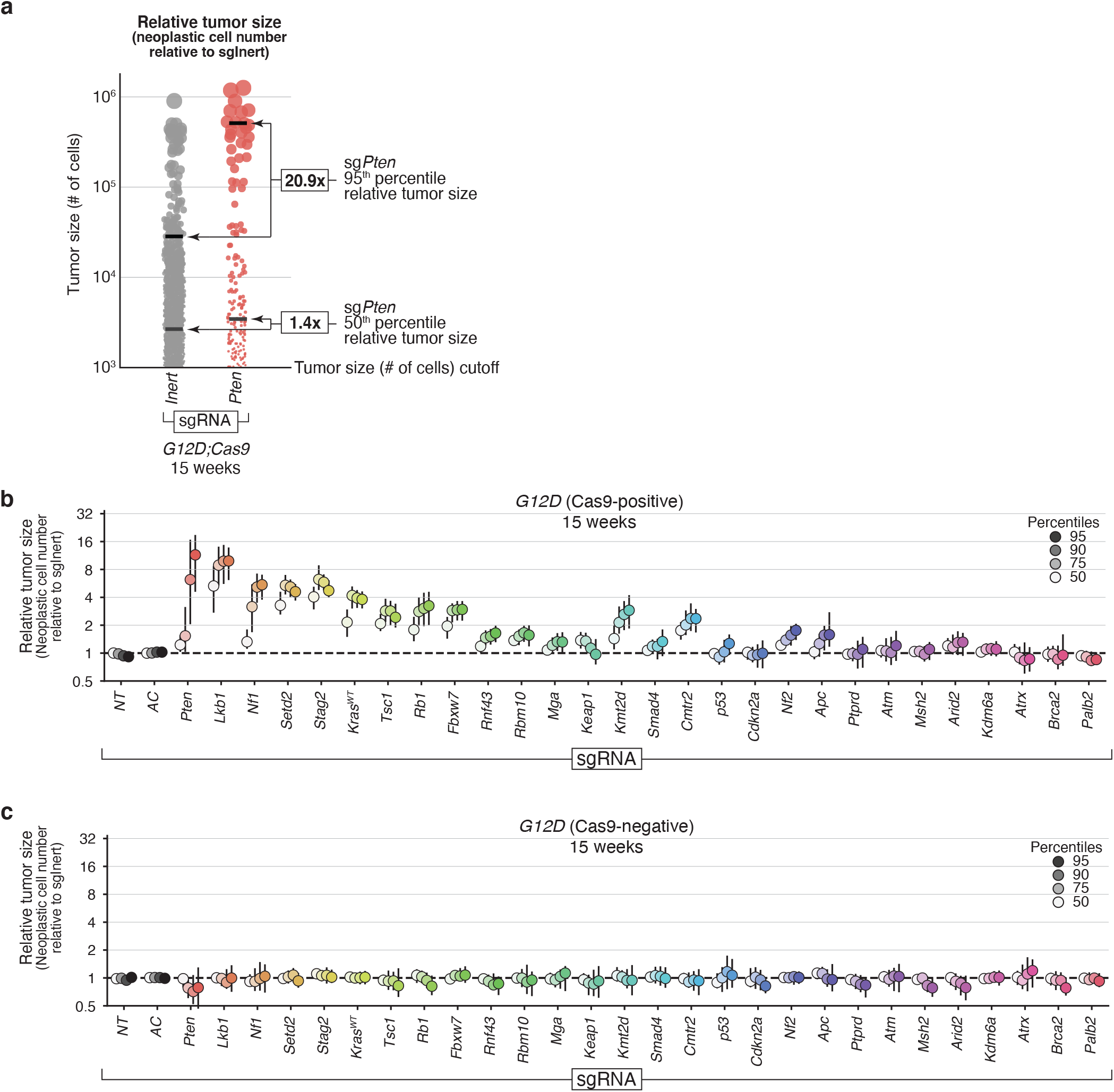
Relative tumor size percentiles enable quantification of effects of Cas9-mediated tumor suppressor inactivation on tumor growth. **a**. Schematic representation of the calculation of relative tumor size (neoplastic cell number relative to sgInert) using simulated sgInert and sg*Pten* tumor size distributions as an example. For all tumors above a defined tumor size (# of neoplastic cells) cutoff (1000 cell cutoff is shown), the size (# of neoplastic cells) of the 50^th^ percentile (median) tumor within the sg*Pten* tumor size distribution divided by the size of the 50^th^ percentile tumor within the sgInert tumor size distribution is used to calculate the sg*Pten* 50^th^ percentile relative tumor size. Relative tumor size can be calculated using the same procedure for any matching percentile (the 95^th^ percentile is also shown) tumor in the sg[*TumorSuppressor*] versus sgInert tumor size distributions. Note that this data from *KrasG12D:Cas9* mice is from a repeat study (corresponding to Group 3 in **Supplementary Fig. 5a,b**) which is distinct from that shown in **Fig. 2a.** **b,c**. Relative size (neoplastic cells) of the tumor at the indicated percentiles of the tumor size distributions for barcoded Lenti-sgRNA/Cre vectors targeting each gene, relative to the size of the sgInert tumor at the same percentile in *G12D;Cas9* mice (**b**) and *G12D* (Cas9-negative) mice (**c**) at 15 weeks after tumor initiation. In the absence of Cas9, all sgRNAs are functionally inert and have no effect on tumor sizes.

**Supplementary Figure 5.**
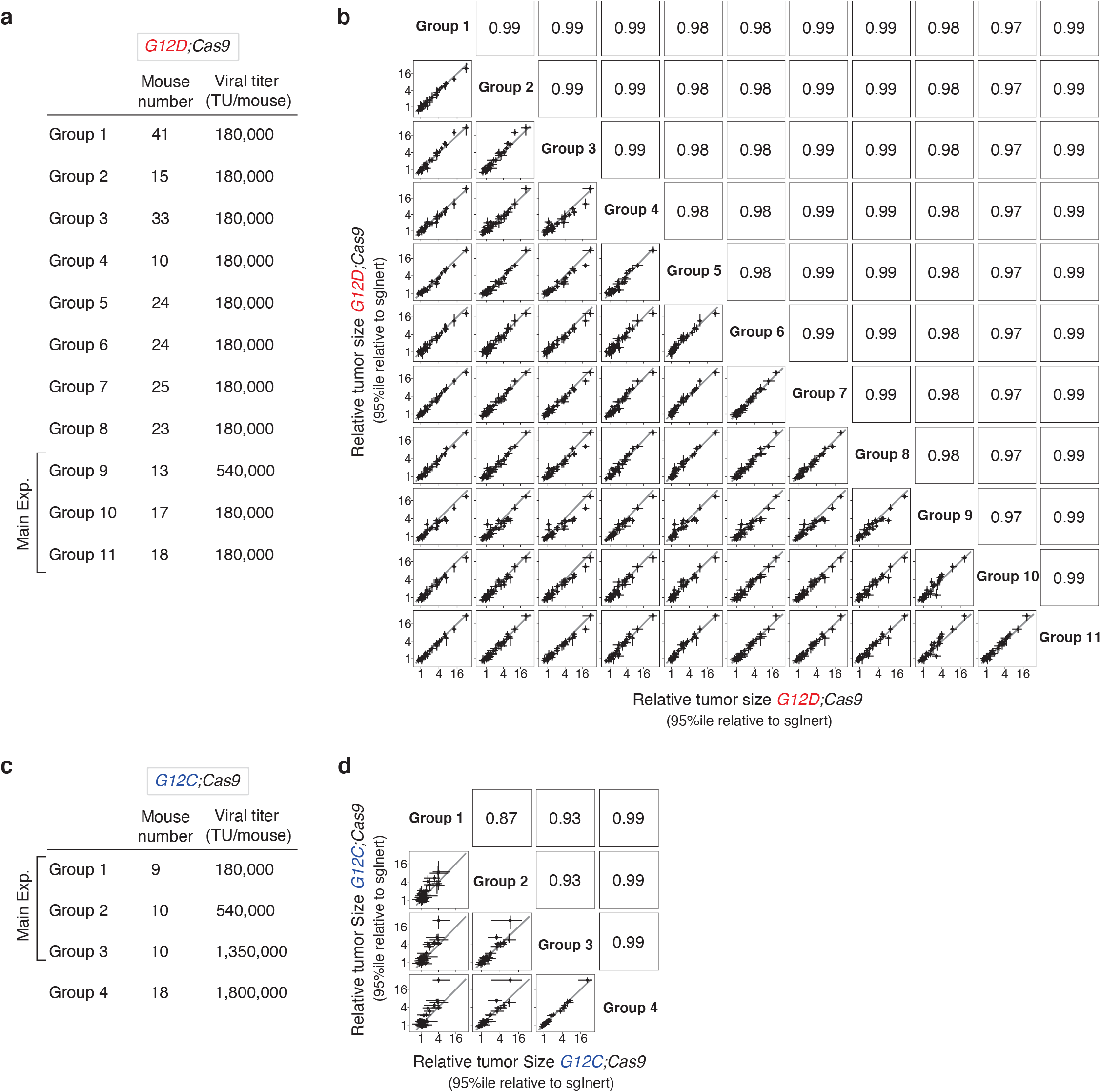
Tumor suppressor effects are highy reproducible across studies. **a,c**. Replicate studies and study groups used to assess the reproducibility of the impact of inactiving tumor suppressors on lung cancer growth in experiments with *G12D;Cas9* (**a**) and *G12C;Cas9* (**c**) mice. Mouse number and viral titers are shown for each group. **b,d**. 95^th^ percentile Relative tumor sizes (relative to sgInert) for each of the 11 *G12D;Cas9* study groups (**b**) and 4 *G12C;Cas9* study groups (**d**). Each point represents the tumors initiated with one Lenti-sgRNA/Cre vector and the bars are the 95^th^ percent confidence intervals. Grey line indicates equal effect. Pearson r is indicated. Note that 3 pairs of groups (1 and 2; 6 and 7; 10 and 11) in panel “**a**” are replicate groups of mice within the same study, while all other listed Groups are from distinct, independent studies initiated at different times.

**Supplementary Figure 6.**
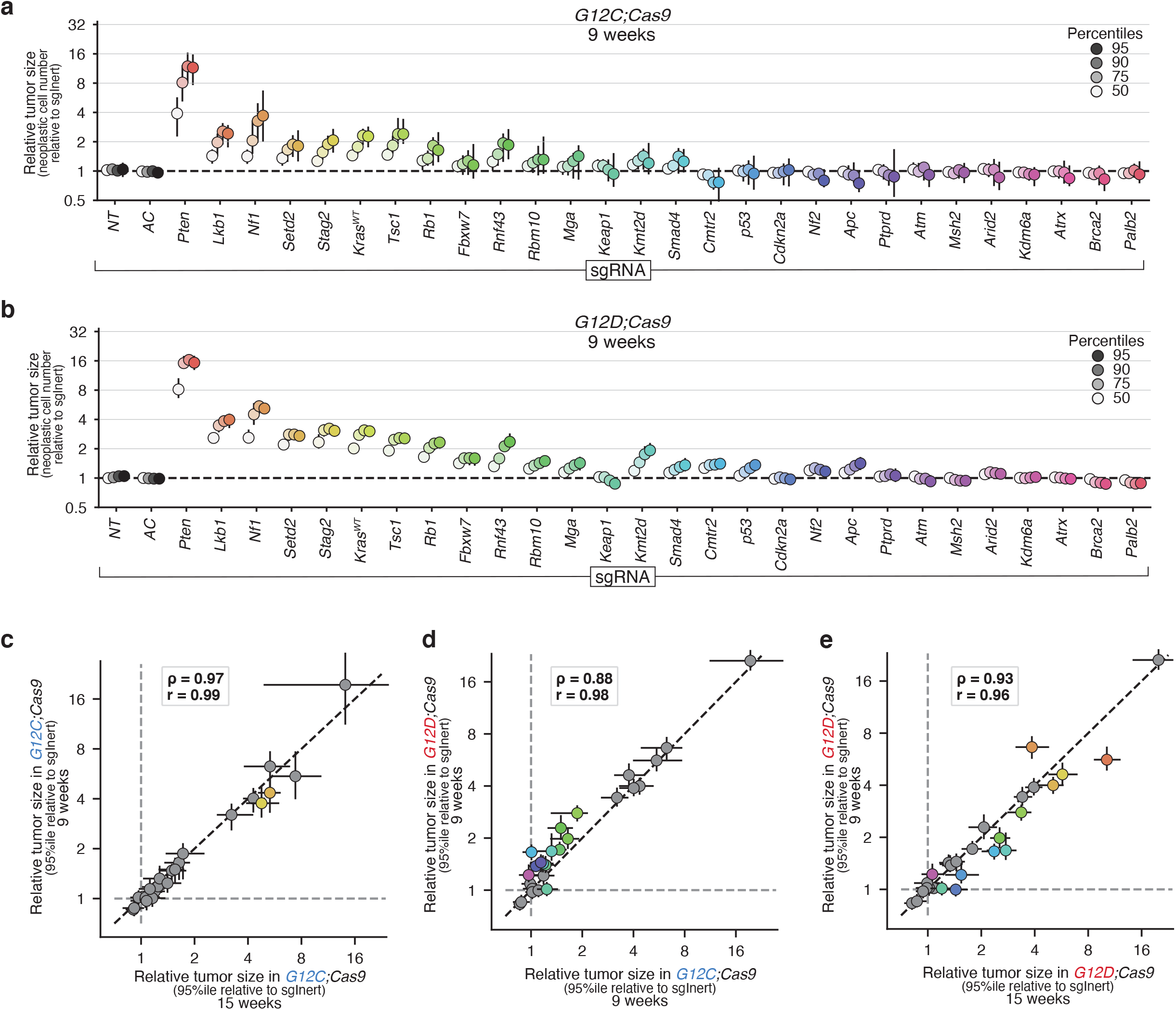
Tumor suppressor effects are reproducible across different timepoints and studies. **a,b**. Relative size (neoplastic cells) of the tumor at the indicated percentiles of the tumor size distributions for barcoded Lenti-sgRNA/Cre vectors targeting each gene, relative to the size of the sgInert tumor at the same percentile, in *G12C;Cas9* mice (**a**) and *G12D;Cas9* mice (**b**) at 9 weeks after tumor initiation. 95% confidence intervals are shown. **c-e**. Relative size of the tumor at the 95^th^ percentile of the tumor size distributions in the indicated genotypes of mice at the indicated times after tumor initiation. Each dot represents the tumors initiated from one Lenti-sgRNA/Cre vector and the bars are the 95^th^ percent confidence intervals. Genes where the 95% CI excluded no effect in both of the two groups are shown in color. Black dotted line indicates equal effect. Spearman rank-order correlation (ρ) and Pearson correlation (r) are indicated.

**Supplementary Figure 7.**
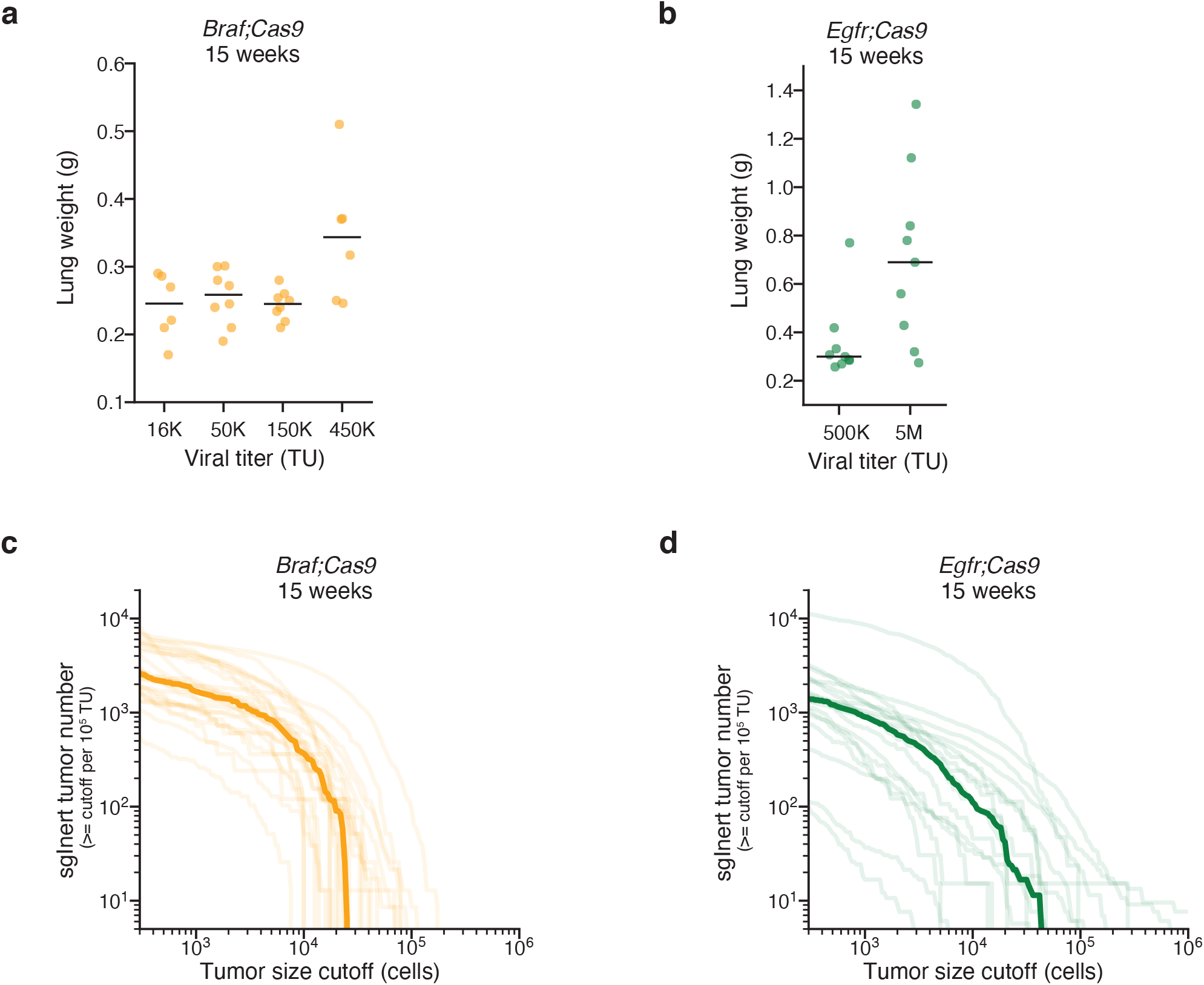
Oncogenic BRAF and EGFR drive the formation of lung tumors with differently shaped size distributions *in vivo*. **a,b**. Lung weights of mice transduced with the indicated titers of Lenti-D2G^28-Pool^/Cre in *Braf;Cas9* (**a**) and *Egfr;Cas9* (**b**) at 15 weeks post-tumor initiation. Each dot represents a mouse and the bar is the median. **c,d**. Number of tumors at or above the tumor size cutoff on the x-axis in *Braf;Cas9* (**c**) and *Egfr;Cas9* (**d**) at 15 weeks post-tumor initiation. Each transparent line represents a mouse and the solid line is the median.

**Supplementary Figure 8.**
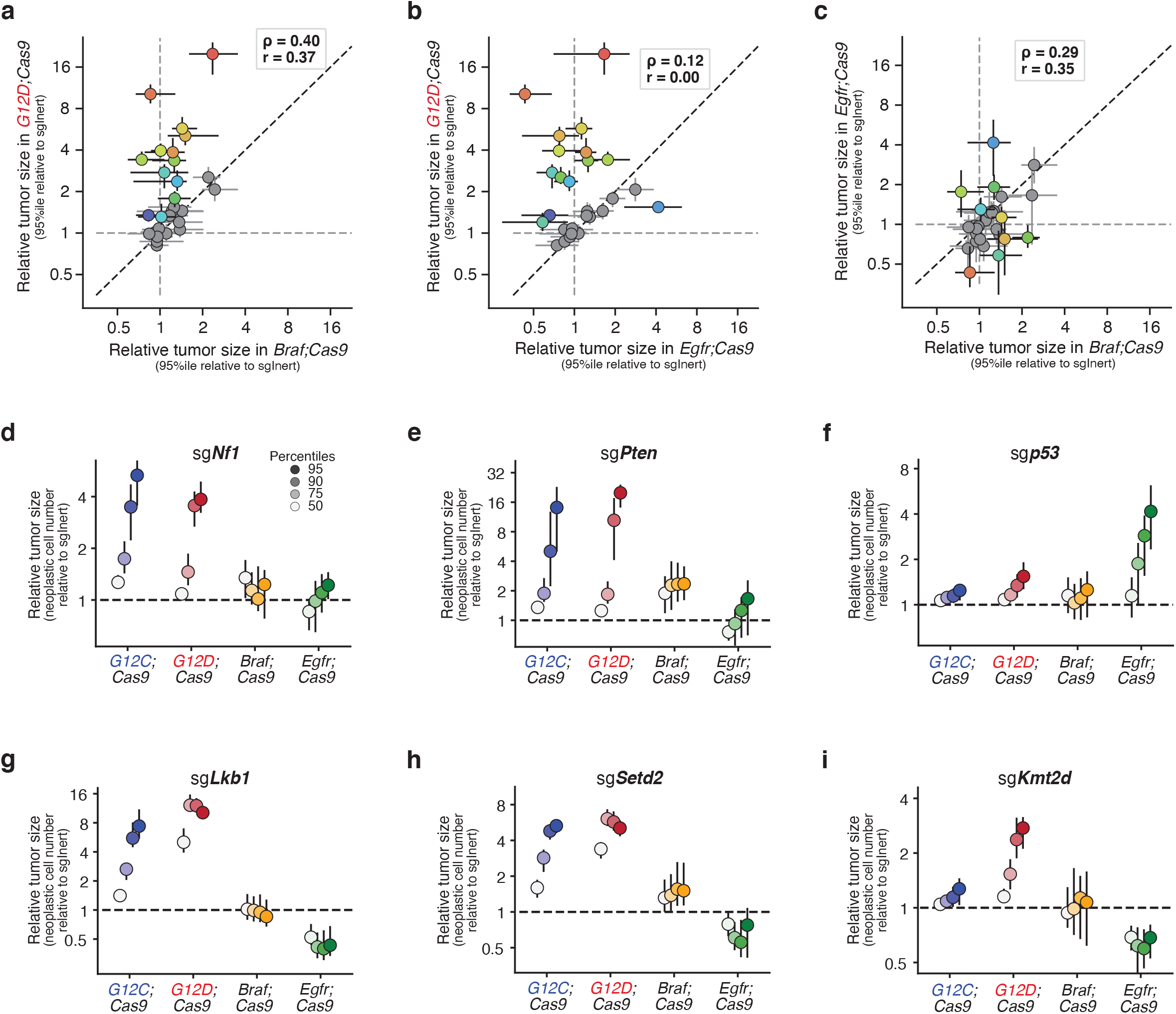
Effect of tumor suppressor inactivation is dependent on oncogenic context. **a-c**. Relative size of the tumor at the 95^th^ percentile of the tumor size distributions in the indicated genotypes of mice at the indicated times after tumor initiation. Each dot represents the tumors initiated from one Lenti-sgRNA/Cre vector and the bars are the 95^th^ percent confidence intervals. Genes where the 95% CI excluded no effect are shown in color. Black dotted line indicates equal effect. Spearman rank-order correlation (ρ) and Pearson correlation (r) are indicated. **d-i**. Relative size of the tumor at the indicated percentiles (legend in **d**) of the tumor size distributions for Lenti-sgRNA/Cre vectors targeting *Nf1* (**d**), *Pten* (**e**), *p53* (**f**), *Lkb1* (**g**), *Setd2* (**h**), and *Kmt2d* (**i**) in tumors in the indicated genotypes of mice.

**Supplementary Figure 9.**
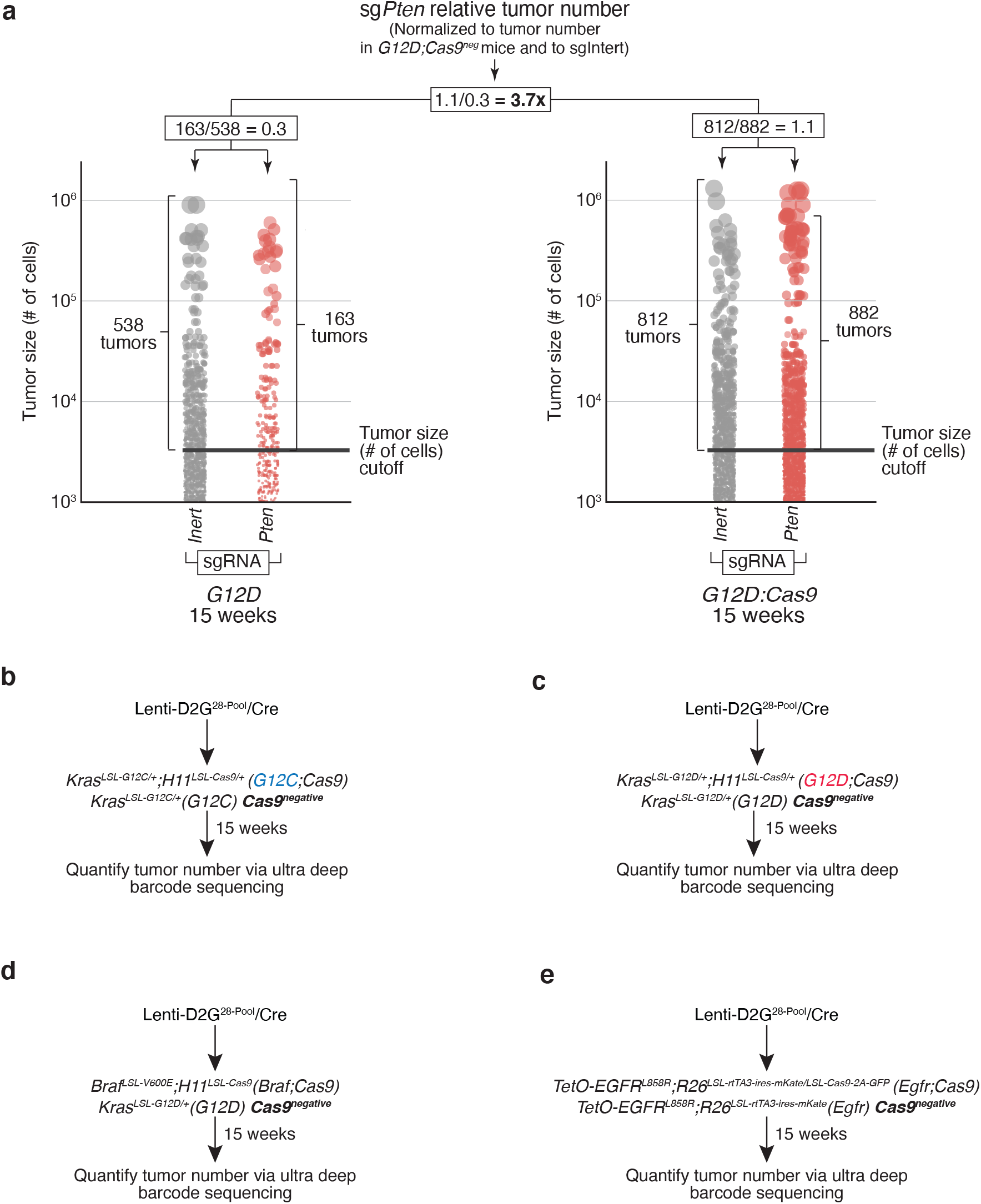
Relative tumor number measurements enable quantification of effects of Cas9-mediated tumor suppressor inactivation on tumor number. **a**. Schematic representation of the calculation of relative tumor number (normalized to tumor number in *Cas9*^*neg*^ mice and to sgIntert) using simulated sgInert and sg*Pten* tumor size distributions as an example. To calculate the sg*Pten* relative tumor number: for all tumors above the tumor size cutoff, the ratio of the number of sg*Pten* tumors to sgInert tumors in the *G12D;Cas9* mice is divided by the ratio of the number of sg*Pten* tumors to sgInert tumors in the *G12D* mice. **b-e**. Description of the mouse genotypes used to calculate relative tumor numbers for *G12C;Cas9* (**b**), *G12D;Cas9* (**c**), *Braf;Cas9* (**d**), and *Egfr;Cas9* (**e**) in **Fig. 5a-d**.

**Supplementary Figure 10.**
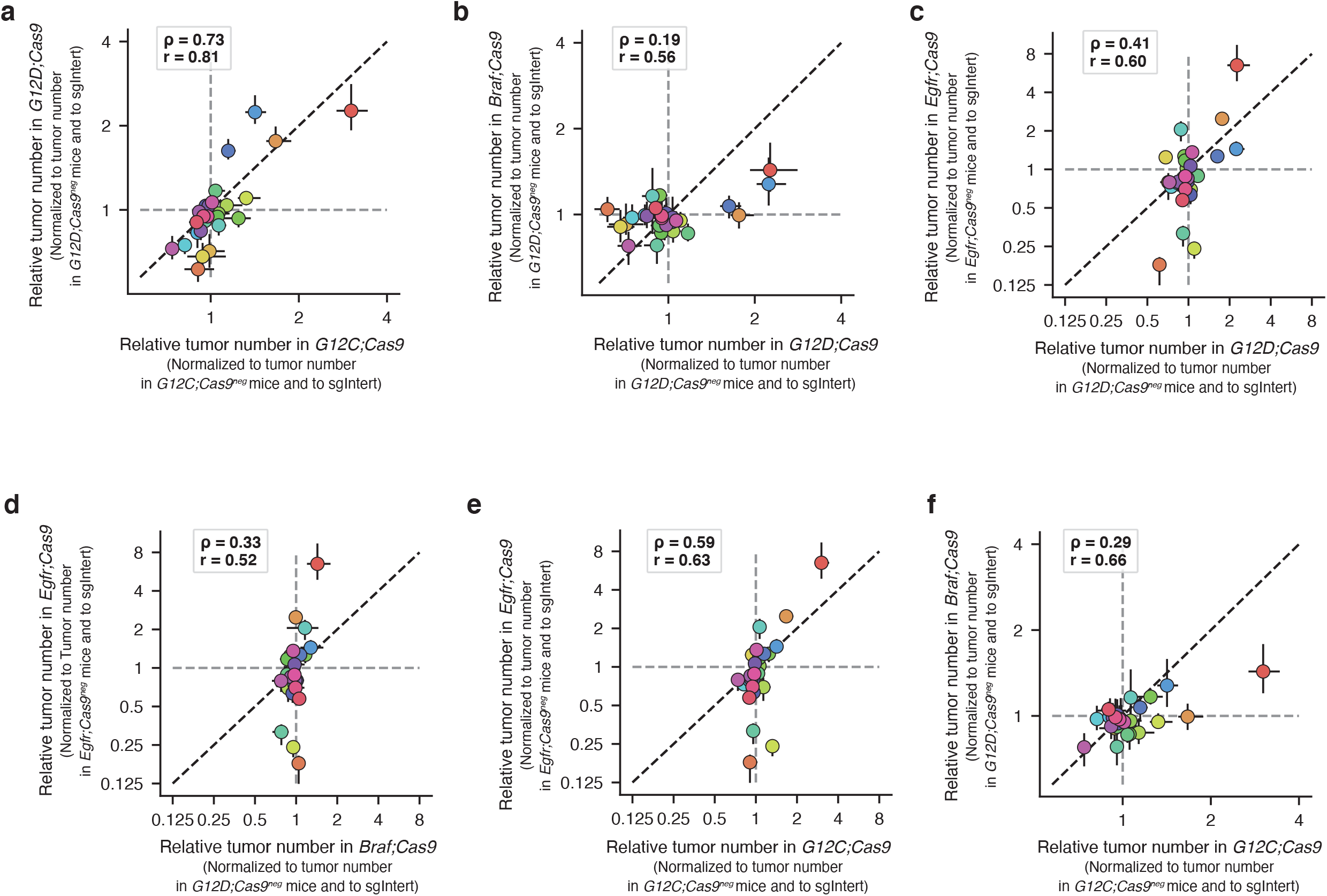
Relative tumor number measurements vary across oncogenes. **a-f**. Relative tumor number in indicated genotypes of mice at 15 weeks after tumor initiation. Each dot represents the tumors initiated from one Lenti-sgRNA/Cre vector and the bars are the 95^th^ percent confidence intervals. The black dotted line indicates equal effect. Spearman rank-order correlation (ρ) and Pearson correlation (r) are indicated.

**Supplementary Figure 11.**
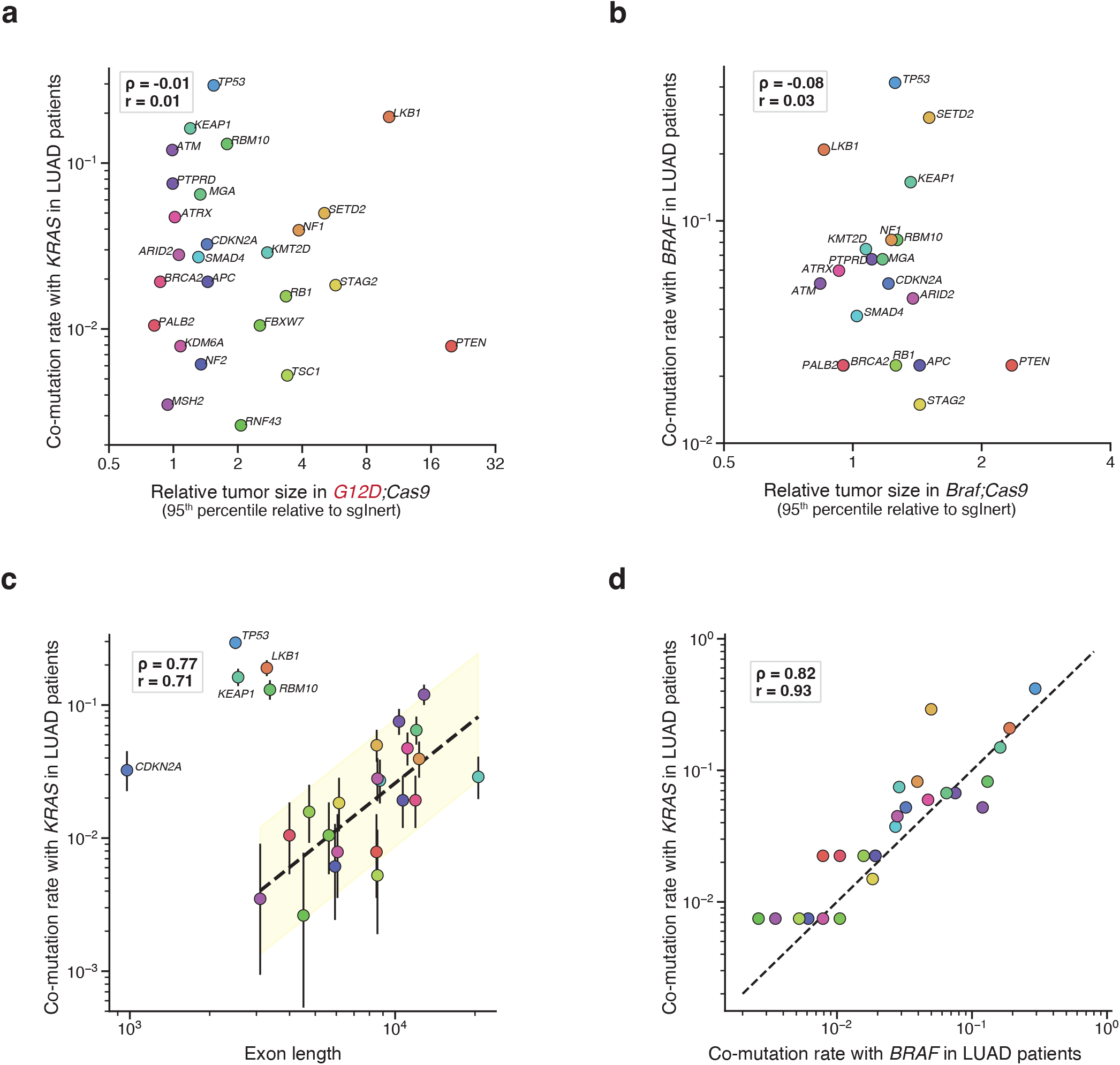
Co-mutation rates in KRAS- and BRAF-driven lung adenocarcinoma are more correlated with exon length than causal effects, suggesting human frequencies in these contexts are driven primarily by passenger mutations. **a,b**. Correlation of relative tumor size at the 95^th^ percentile to co-mutation rate of each gene tested in our model with *KRAS* (**a**) and *BRAF* (**b**) in LUAD patients. *CMTR2* was the only gene tested in our model that was not present in the MSK-IMPACT468 panel and therefore not included here. Spearman rank-order correlation (ρ) and Pearson correlation (r) are indicated. **c**. Correlation of exon length to co-mutation rate of each gene tested in our model with *KRAS* in LUAD patients. Error bars show the 95% binomial confidence interval.*TP53, LKB1, KEAP1, RBM10*, and *CDKN2A* were determined to be outliers. After removal of outliers: dotted line shows linear fit to the log-transformed exon length and log-transformed co-mutation rate; yellow region highlights 3x on either size of the fit, spearman rank-order correlation (ρ) and Pearson correlation (r) are indicated. **d**. Correlation of co-mutation rate wth *BRAF* in LUAD with co-mutation rate with *KRAS* in LUAD patients. Spearman rank-order correlation (ρ) and Pearson correlation (r) are indicated.

**Supplementary Figure 12.**
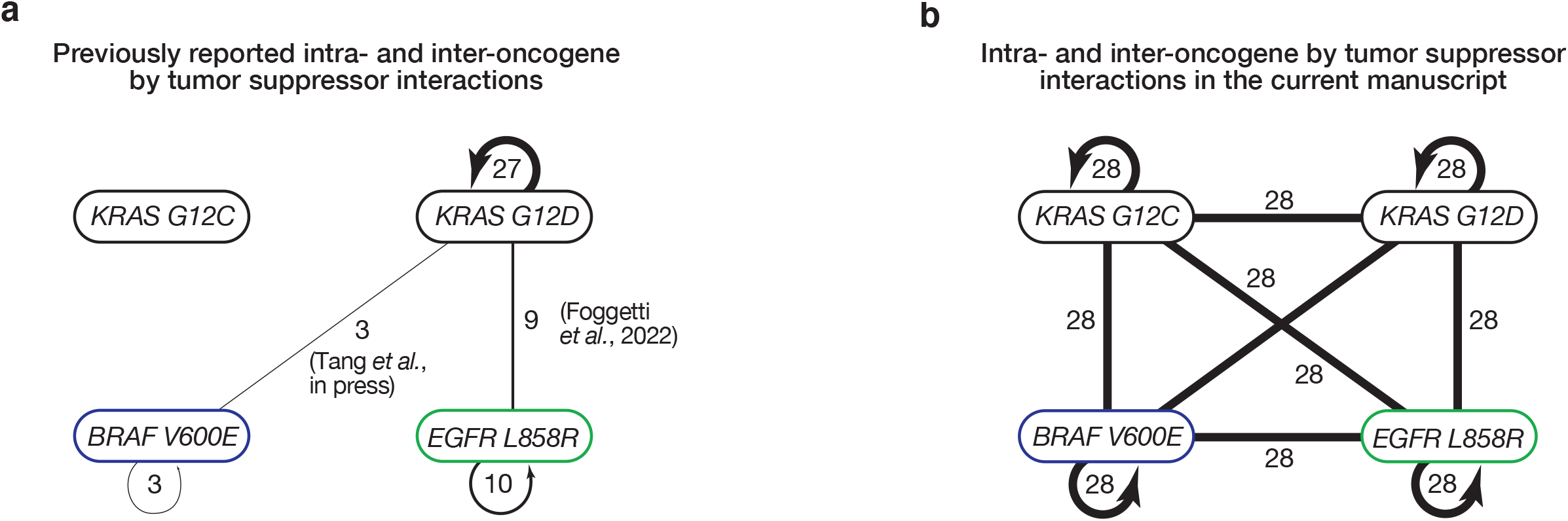
Extent of novel intra- and inter-oncogene comparisons of tumor suppressor effects described in the current manuscript. **a**. Previous data investigating the impact of inactivating putaive tumor suppressor genes on tumor growth within and across oncogenic contexts in quantitative *in vivo* models. **b**. Data from the present study investigating the impact of inactivating putaive tumor suppressor genes on tumor growth within and across oncogenic contexts.

